# Compression of morbidity by interventions that steepen the survival curve

**DOI:** 10.1101/2023.10.04.560871

**Authors:** Yifan Yang, Avi Mayo, Tomer Levy, Naveh Raz, Dan Jarosz, Uri Alon

**Affiliations:** Department of Molecular Cell Biology, Weizmann Institute of Science, Rehovot, Israel; Stanford University School of Medicine, Stanford, CA, USA

## Abstract

Longevity research aims to extend the period of healthy life, known as the healthspan, and to minimize the duration of disability and morbidity, known as the sickspan. Most longevity interventions in model organisms extend healthspan, but it is not known whether they extend healthspan relative to the lifespan - that is, whether they compress relative sickspan. Here, we present a theory that predicts which interventions effectively compress relative sickspan. The theory is based on the shape of the survival curve - the fraction of organisms surviving as a function of age. Interventions that extend mean lifespan while preserving the shape of the survival curve, known as scaling, are predicted to extend the sickspan proportionally, without compressing it. Such interventions include caloric restriction and many other longevity interventions. Conversely, a small subset of interventions that extend lifespan and steepen the shape of the survival curve, are predicted to compress the relative sickspan. We explain this based on the saturating removal mathematical model of aging, and present evidence from longitudinal health data in mice, *Caenorhabditis elegans* and *Drosophila melanogaster*. We discuss longevity interventions in mice that steepen the survival curves, including senolytics, ketogenic diet, and agents that reduce glucose spikes and protect blood vessels, as potential candidates for compressing the sickspan. We apply the theory to combinations of longevity interventions and discuss human healthspan data. This approach offers potential strategies for compressing morbidity and extending healthspan.

## Introduction

The lifespan of an individual typically begins with a period of general health, called the healthspan, followed by a period of intermittent or continuous morbidity and disability called the sickspan ^1–3^. The sickspan burdens human quality of life and is a growing driver of economic expenditure across societies ^4–6^.

Understanding what sets lifespan, and how to extend it, is a central goal of longevity research. There are several possible scenarios of lifespan extension regarding sickspan ^4^. In principle, a longevity intervention could expand morbidity, and in the extreme case extend the sickspan exclusively, which is clearly undesirable. Alternatively, both healthspan and sickspan could be simply ‘stretched’ in proportion to lifespan. Finally, as proposed in the field of gerontology ^1–3^, one could ideally aim to compress morbidity. Sickspan can be compressed either in absolute terms, as advanced by Fries ^7^, or relative to lifespan ^8^, as in the dynamic equilibrium concept by Manton in which both healthspan and lifespan are extended by similar absolute amounts ^9^. Generally, optimal aging trajectories restrict morbidity to a brief period before death ^3,7^.

Research has identified diverse interventions that can extend lifespan in model organisms including yeast, nematodes, flies and mice ^10–14^. These include dietary restriction ^15,16^, perturbations of the IGF1 pathway ^17^, mTOR inhibition by drugs such as rapamycin ^18,19^, senolytic treatments ^20–22^, *in vivo* partial reprogramming ^23,24^ and additional classes of drugs including antioxidants and diabetic drugs^25^.

These longevity interventions are developed to delay morbidity in model organisms - the treated animals are usually reported to have better health parameters than equally aged untreated animals as measured in a single time point ^11^. However, whether they compress the sickspan in absolute or relative terms has not generally been reported. Data from *Caenorhabditis elegans* suggests that many interventions that extend lifespan extend sickspan proportionally ^26,27^, and thus do not compress morbidity in absolute or relative terms. A similar effect has been seen in medflies ^28^ . Notably, however, intervention studies that longitudinally follow health in individual organisms are rare, especially in mammals, and thus the question of sickspan relative to lifespan is understudied.

It is thus of interest to identify which longevity interventions are likely to compress sickspan, and which instead have the unintended consequence of extending it. Identifying factors that affect the relative healthspan and sickspan in model organisms would improve our basic understanding of the biology of aging and age-related morbidity and inform the search for healthspan-extending therapies.

Here we present a theory that can predict which longevity interventions compress sickspan. The theory is based on the survival curve, the fraction of organisms that survive to a given age ^29^. Many longevity interventions that extend median lifespan preserve the shape of this curve, a phenomenon called scaling (Fig. 1b) ^26,30–32^. We predict that such interventions extend sickspan proportionally. In contrast, a small number of interventions extend median lifespan and steepen the survival curve, making its shape more rectangular. We predict that these interventions compress the sickspan (with certain exceptions discussed below).

**Figure 1.**
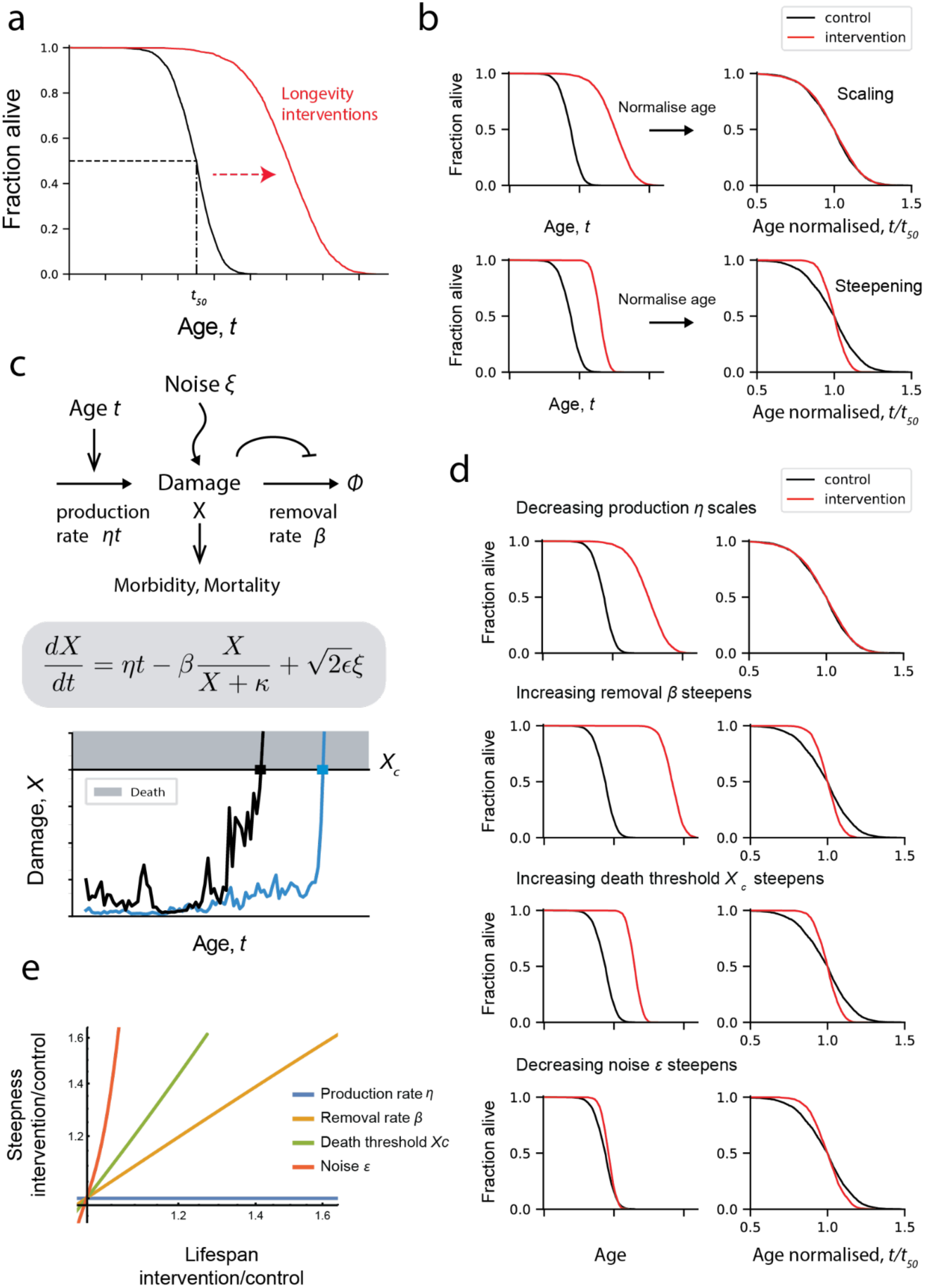
Survival curves in the saturating removal model scale when interventions affect damage production, and steepen when interventions affect other parameters. a, Survival curves describe the fraction of organisms surviving to each age; median age *t50* is extended by longevity interventions. b, Scaling is defined by survival curves collapsing on the same curve when time is re- normalized by their averages ^30^. Steepening occurs when the intervention’s survival curve has higher steepness than control. c, The saturating removal model posits a damage *X* that drives aging whose production rises with age and whose removal saturates, with noise. Simulations (black and blue lines) show stochastic trajectories of 2 individuals that cross the death threshold Xc at different times. d, Effect of a 2-fold change in each model parameter on survival curve shape e, compared to baseline, effect of parameter changes on *t50* and steepness defined as median divided by interquartile range. Baseline parameters for mice are *η*=2.3x10^-4^/day^2^, *β*=0.15 /day, *κ*=0.5, *ε*=0.16 /day, *Xc*=17 ^32^.

Here we show that this theory arises naturally from a recent mathematical model of aging called the saturating removal model that was calibrated in mice and single celled organisms ^32–35^. We provide evidence from invertebrate and mice data that steepening interventions compress relative sickspan. We discuss specific mice longevity interventions that steepen the survival curve as candidate interventions to compress sickspan, and their potential mechanisms, and apply the theory to combinations of longevity interventions and to human data.

## Results

### The saturating removal model links the shape of survival curve with the parameter of damage dynamics

Longevity-enhancing interventions shift the survival curve to greater ages, increasing the median survival age *t50* (Fig. 1a). They often preserve the curve shape, a phenomenon called scaling ^26,30,31^. Interventions are said to be scaling if their survival curves collapse onto the control survival curve when age is normalized by the median lifespan (Fig. 1b, upper panel, we use accelerated failure time statistical tests for scaling, Methods). Scaling interventions increase median and maximal lifespan by the same factor. Scaling was discovered by Strousturp and colleagues in *C. elegans* ^30^, and later described in budding yeast ^31^.

For example, median C elegance lifespan increases at low temperatures, but survival curves scale to a good approximation. Caloric restriction also extends lifespan ^15,16,36^, as reviewed by Fontana, Partridge and Longo ^16^, but survival curves approximately scale.

Other interventions increase median lifespan but steepen the survival curve shape (Fig 1b, lower panel) ^30,37^ . A useful metric for steepness is the median age divided by the interquartile range of survival ^29^. In the following we use an accelerated failure time method, following previous works ^30,38^, to determine whether a curve scales or not (Methods).

Interventions that extend lifespan and scale the survival curve seem to stretch the time axis as if time is slowed down. They may therefore be considered intuitively to slow down the overall pace of aging. We hypothesize that they should accordingly slow down the processes responsible for age-related morbidity and functional decline. The result is a proportional scaling of the sickspan and healthspan. Relative sickspan, that is the sickspan divided by lifespan, should remain unchanged by the intervention.

To quantitatively test this intuition, we use the saturating-removal (SR) mathematical model for aging dynamics ^32,34,35^. (Fig. 1c). The SR model describes the dynamics of damage that is posited to be causal for both morbidity and mortality. The central simplifying assumption of the model is that there is a dominant form of damage that is upstream of the vast range of age-related changes in the organism. The model is agnostic to the molecular identity of this driving damage and can thus apply to different organisms despite the difference in their mechanisms of aging. The SR model is a stochastic differential equation ^39^ whose mathematical form was deduced from dynamic data on mouse senescent cells. Its predictions were tested experimentally in mice ^32^ and in longitudinal damage measurements in microorganisms ^35^. The model was further validated by comparison to quantitative datasets on age related diseases in humans ^34^, parabiosis in mice ^33^ and midlife longevity interventions in drosophila ^32^.

The key simplifying assumption in the model is the existence of a driving damage denoted *X* which is upstream of morbidity and mortality. The model is agnostic to the exact nature of *X* and thus can apply to different organisms. The driving damage rises with age and fluctuates around this rising trend on a timescale that is fast compared to the lifespan. The model is a stochastic differential equation for *X* that accounts for production, removal and noise: *dX*/*dt* = production - removal + noise.

The driving damage *X* is produced in the model by damage-producing units that are not removed and accumulate linearly with age (Fig. 1c). The rate of production is therefore proportional to age, namely *production = ηt*. The driving damage is removed by removal processes that saturate at high damage levels. This is modeled as Michaelis-Menten-like saturation, *removal = βX*/(*κ*+*X*), where *β* is the maximal removal rate and *κ* the halfway saturation point. The noise is modeled as white noise with amplitude *ε*. The SR model equation is shown in Fig. 1c.

Death is modeled to occur when driving damage *X* crosses a mortality threshold *Xc* (Fig. 1c). Simulating the model shows that each individual has a stochastic trajectory of damage *X* (Fig. 1c). Each individual therefore crosses the death threshold *Xc* at a different time. The risk of death in the model rises exponentially with age and slows at very old ages ^32^ - this is the Gompertz law found in many species, including humans. The survival curves are sigmoidal in shape (well-described by Weibull distributions) and resemble empirical survival curves ^32^.

A key feature of the SR model is its ability to provide scaling of survival curves ^32^. Changes in the rate of damage production, namely the parameter *η*, shift median survival but preserve the shape of the curve (Fig. 1d), providing near-perfect scaling.

Changing any of the other model parameters, such as removal rate *β*, affects not only lifespan but also the shape of the survival curve (Fig. 1d). Increasing damage removal *β* shifts the survival curve to the right and steepens it. Increasing death threshold *Xc* mildly extends median lifespan and steepens the curve by preventing early deaths. Decreasing the noise amplitude *ε* causes a small increase in lifespan and a large increase in steepness - less noise leads to more deterministic death times. These effects are shown on a steepness-longevity plot in Fig. 1e, and separately for steepness and longevity in Supp. Fig. 1.

The reason that these parameters all affect steepness is that the shape of the survival curve in the model is essentially determined by a single dimensionless parameter *βXc*/*ε*. Note that production rate *η* does not appear in this dimensionless group. It affects lifespan but not the shape of the curve and so changes in *η* preserve scaling.

### Interventions that steepen the survival curve compress the relative sickspan in the saturating removal model

Our main interest in this study is sickspan. Hence, we add morbidity to the model to study sickspan and healthspan. Age-related morbidity is modeled to occur when *X* crosses a morbidity threshold *Xd* (Fig. 2a). This simple model captures the incidence of hundreds of age-related diseases in humans as shown by Katzir et al ^34^. Each disease has its own threshold. Here we consider sickspan as an aggregate phenomenon that occurs when a threshold is crossed that is an average of the different disease thresholds weighted by their prevalence. The probability of crossing *Xd* rises exponentially with age until it slows at very old ages.

**Figure 2.**
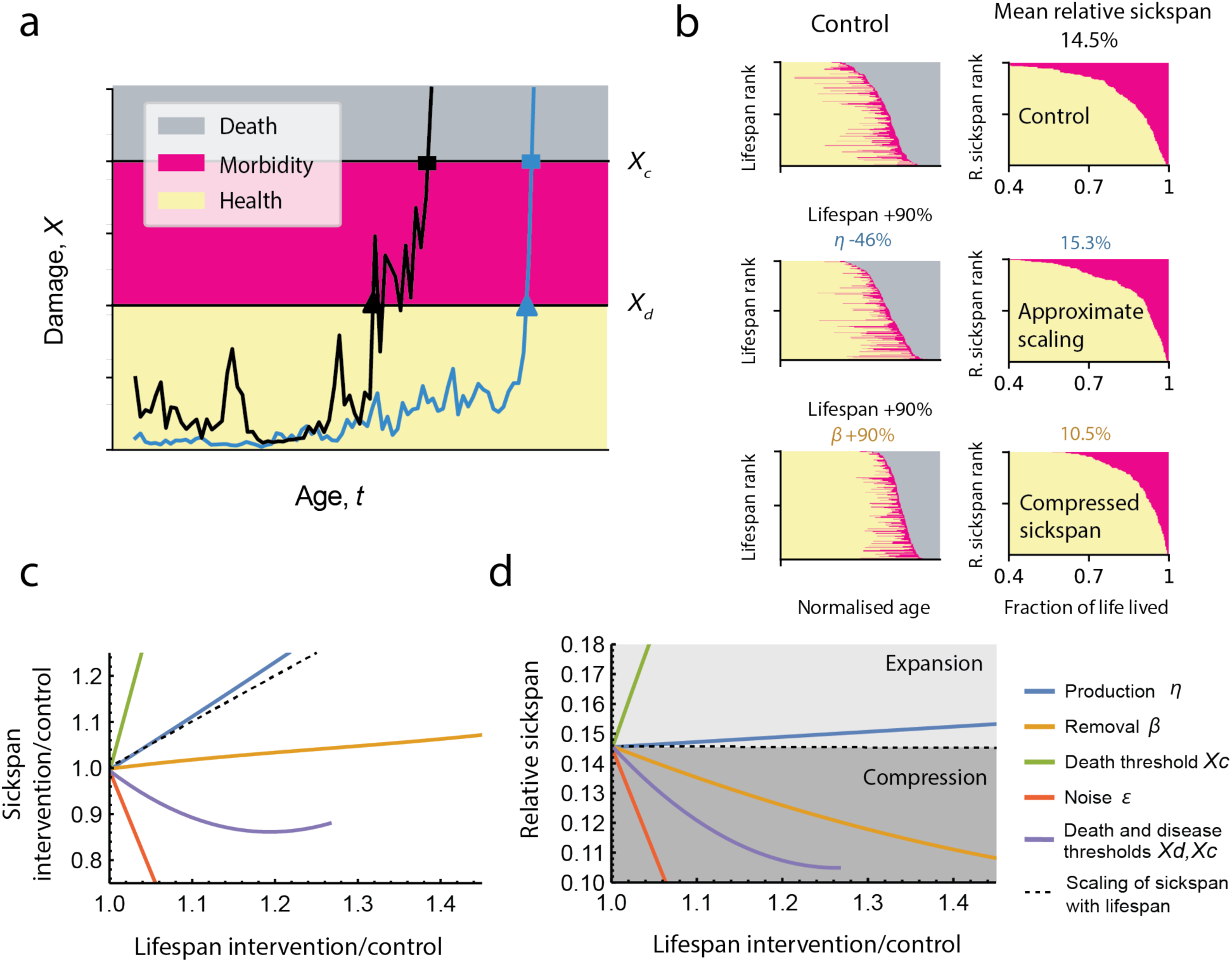
Interventions that scale survival curves also scale the sickspan, whereas interventions that steepen survival curves compress relative sickspan. a, Trajectories of damage *X* for two simulated individuals. Sickspan is the period after first crossing the disease threshold *Xd* and before crossing the death threshold *Xc*. **b,** Simulation of untreated individuals and two treatments that extend median lifespan by the same amount: reduced production *η* by 46% or increased removal *β* by 90%. Simulations of 200 individuals are shown in two ways: left, ranked by lifespan versus age (survival curve); right, ranked by relative sickspan as function of fraction of life lived. Red indicates the sickspan. **c,** Absolute sickspan (intervention/control) as a function of median lifespan for changes in different model parameters. **d,** Relative sickspan as a function of model parameters. Baseline parameters are as in Fig. 1 with *Xd*=12.

We thus consider here a definition of sickspan as a continuous end-of-life period that begins upon the first crossing of *Xd*. This aligns with current definitions in model organisms of an end of life period of morbidity when severe health deficits cross a threshold ^26,40–42^. An alternative definition would allow for sickspan to be intermittent when *X* drops below *Xd* for sufficient time. Such a definition requires additional parameters and assumptions and we leave it for future study.

We tested the effect of changing model parameters on the relative sickspan. We find that changes in production *η* that scale the survival curve also approximately scale the relative sickspan - and in fact slightly prolong it (Fig. 2b-d).

In contrast, sickspan is compressed by most changes that steepen the survival curve. Increasing removal rate *β* compresses the relative sickspan (Fig. 2b-d), as does a reduction in noise *ε* or increasing both disease and death thresholds together (Fig. 2c,d). In these interventions, longevity is increased primarily by extending the healthspan.

There is one exception to the rule that interventions that steepen survival curves compress relative sickspan. This occurs when the death threshold *Xc* rises but the disease threshold *Xd* does not (Fig. 2c,d, green lines). This might apply to human data when medical care extends life in sick individuals, without affecting age of onset of sickspan. We discuss human mortality in the last section.

In summary, longevity interventions have differential effects on relative sickspan depending on which parameters they affect in the model. Changes in damage production rate *η* - the only changes that scale the survival curve - increase absolute sickspan and mildly increase relative sickspan. In contrast, interventions that extend lifespan and steepen the survival curve generally compress the relative sickspan. They extend lifespan primarily by extending healthspan.

### Steepening interventions compress relative sickspan in invertebrates and mice

To test whether steepening of the survival curve indeed implies a compressed sickspan, we analyzed data from experiments which followed health longitudinally as organisms aged under different interventions. These experiments were performed in mice, *Drosophila melanogaster* and *Caenorhabditis elegans*. In all cases we used the authors’ definition of sickspan.

Data on longitudinal health in mice is currently limited, although it is expected to be available from several ongoing experiments ^43–45^. We analyzed preliminary data from Luciano et al ^42^ which followed mouse frailty index longitudinally in different longevity interventions. The experiment included dietary interventions (20% and 40% calorie restriction and two intermittent fasting schedules) in two strains (Diversity Outbred and C57BL/6). Sickspan was defined by the authors as periods of life with equal or more than 4 age-related deficits in the mouse frailty index. We evaluated the survival curve steepness relative to control for these experiments and compared it to the relative sickspan (mean absolute sickspan divided by mean lifespan, relative to control). Steepness was defined by the median lifespan divided by the interquartile range (IQR) of lifespan ^29^. We calculate confidence intervals of both relative sickspan and steepness by bootstrap resampling of individuals.

We find that relative sickspan decreased with steepness as predicted (Fig. 3a, Supp. Table 1). Scaling interventions (steepness intervention/control of 1) showed approximately scaled sickspan (Fig. 3a). These findings agree with the SR model predictions. The relationship is robust to the definition of sickspan (Supp. Fig. 2).

**Figure 3.**
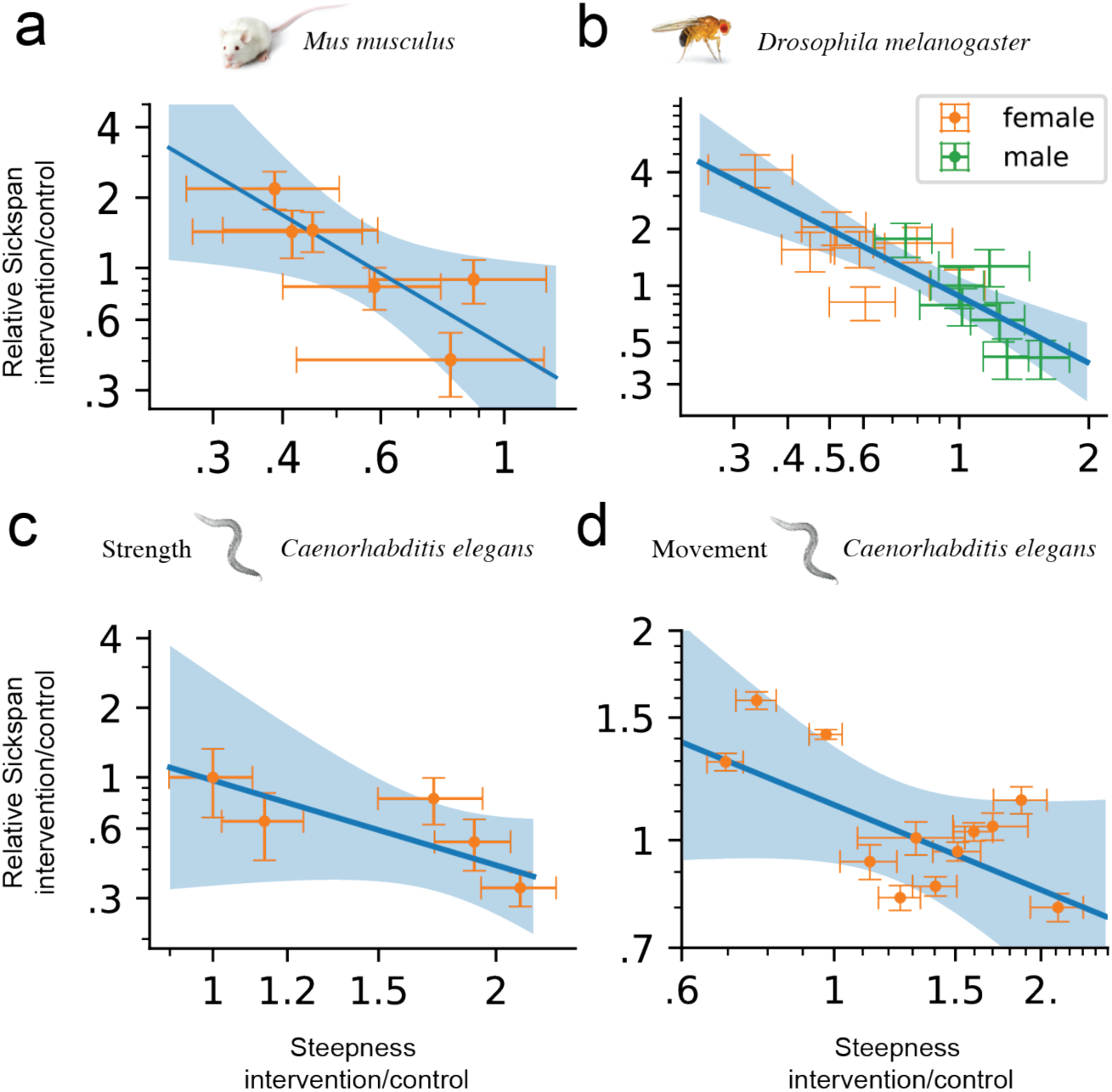
Relative sickspan is compressed by interventions that steepen the survival curves in model organisms, as predicted. Mean absolute sickspan divided by mean lifespan relative to control is plotted versus steepness of survival curve relative to control. **a**, Mice under dietary longevity interventions - caloric restriction and intermittent fasting - in two strains from Luciano et al ^42^. Sickspan was defined using mouse frailty index. See Supp. Table 1 for intervention details. **b**, Nutritional longevity interventions in two strains of *D. melanogaster*, where sickspan was defined by deficits in climbing, jumping and flying in response to a mechanical challenge. Data from Gaitanidis et al ^40^. See Supp. Table 2 for intervention details. **c**, Genetic longevity mutants in *C. elegans*, where sickspan was defined by loss of muscle power measured by piezoelectric challenge. Data from Statzer et al ^26^. See Supp. Table 3 for strain details. **d**, Genetic, dietary and environmental factors in *C. elegans*, where sickspan was measured by secession of spontaneous movements. Data from Stazer et al ^26^ and Oswal et al ^41^. Confidence interventions are calculated using bootstrap resampling of individuals in each cohort. See Supp. Table 4 for strain and intervention details.

We next considered longitudinal experiments in flies. Carey and colleagues pioneered the use of longitudinal life-history data such as fecundity and mortality to test evolutionary theories of aging ^46–48^ using medflies (*Ceratitis capitata*). More recently, Gaitanidis et al ^40^ measured longitudinal healthspan data in *Drosophila* by quantifying age-related declines of motor functions. The authors assayed escape performance in response to gently banging food vials on the counter, i.e. “startle assays”.

Each fly was scored for its jumping, climbing and flying responses to generate a health score for each individual by summing the number of functional deficits. Sickspan was defined as the period where an individual has a score below a threshold value. The experiment was repeated for 5 dietary interventions (including protein restriction, superfood and curcumin) in males and females in two fly strains with different lifespans (Lausanne, Oregon).

We evaluated the survival curve steepness relative to control for these experiments, and compared it to the relative sickspan. Relative sickspan decreased with steepness, as predicted. Scaling interventions (steepness/control of 1) showed approximately scaled sickspan (Fig. 3b, Supp. Table 2). These findings agree with the SR model predictions.

We also considered two *C. elegans* studies. Statzer et al used a piezoelectric system to measure muscle power cross-sectionally throughout the lifespan ^26^, for five different longevity-affecting mutants: daf-2(e1368), daf-2(e1370), glp-1 and eat-2, and define sickspan by the period of life when muscle power is below 50% of wildtype. Analysis of this data shows that relative sickspan decreases with steepness (Fig. 3c, Supp. Table 3).

Another method to define healthspan used the period of spontaneous movements to define the healthspan ^41^. This method was used by both Oswal et al ^41^ and Statzer et al ^26^, for several longevity interventions including temperature, food deprivation, food inactivation by UV, daf-2 (e1368 & e1370), eat-2, glp-1 (both studies), nuo-6 (see Supp. Table 4 for details of all the intervention and control cohorts). The analysis showed that relative sickspan defined by voluntary movement also decreases with survival curve steepness, as predicted (Fig. 3d).

We conclude that in invertebrates and mice, steepening interventions compress the relative sickspan, whereas scaling interventions approximately scale the sickspan.

### Several interventions in mice are predicted to compress sickspan

Whereas longitudinal health data is currently rare, survival curve data of mice under longevity interventions is routinely reported in the literature ^49^. We therefore asked which mouse interventions show scaled survival curves, and which steepen the survival curve ^37,38^.

We explored the 42 nutritional and pharmacological interventions reported by the National Institute on Aging (NIA) Interventions Testing Program (ITP) ^50^ (Fig. 4a, Supp. Data 1, Supp. Fig. 3-5,). The ITP experiments are standardized lifespan experiments done in three parallel sites with genetically heterogeneous mice and relatively large cohorts ^49^. We also analyzed intervention experiments reported in 32 additional publications, including those with transgenic mice (Fig. 4b, Supp. Data 2, Supp. Fig. 6).

**Figure 4.**
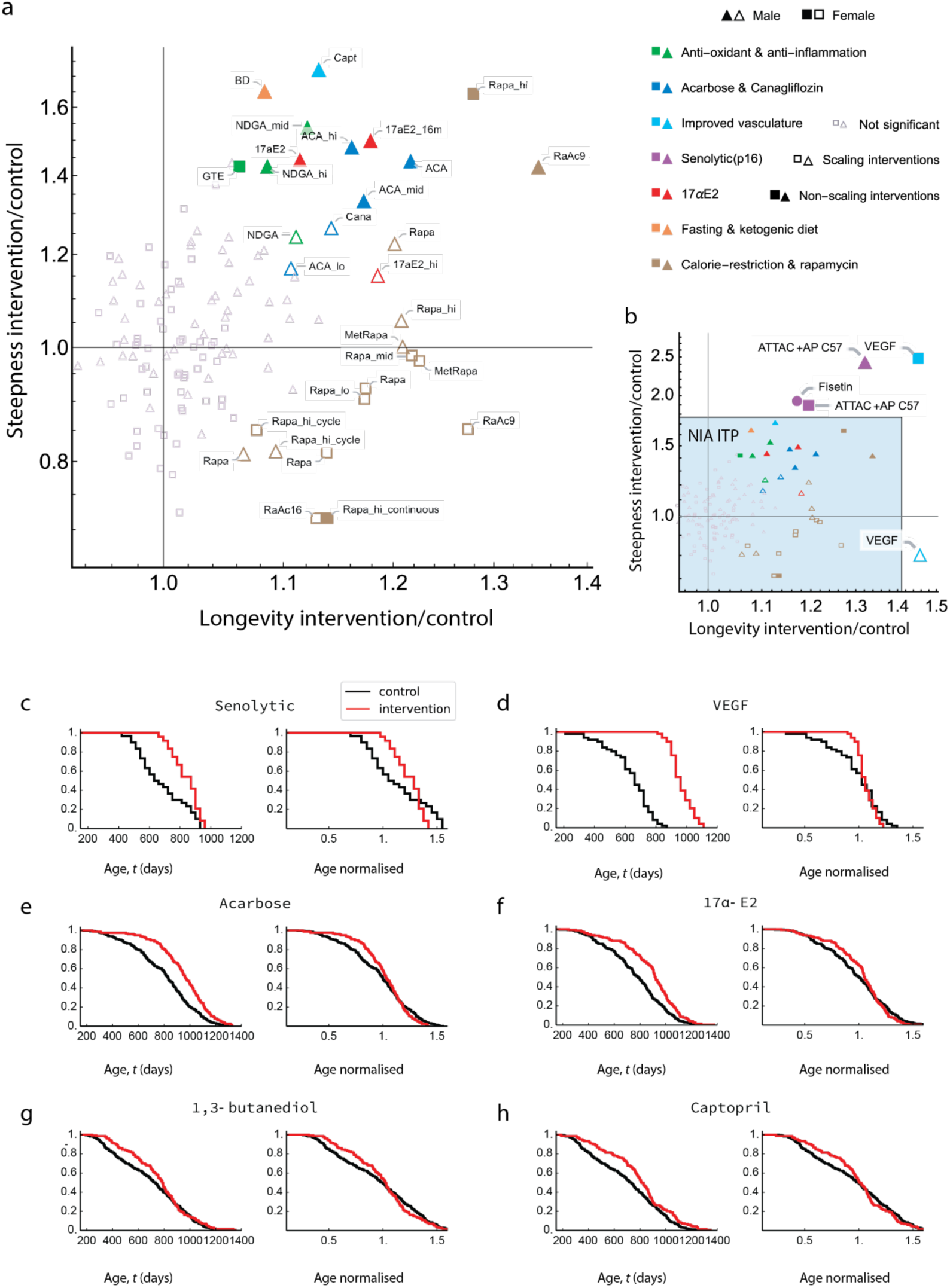
Lifespan extension interventions in mice that steepen the survival curve. **a**, Longevity and steepness relative to control in all interventions from the NIA Interventions Testing Program (NIA ITP) ^50^. Square and triangle symbols denote female and male mice respectively. Small light symbols indicate interventions whose effects are not statistically significant (Hotelling’s t-squared test, α=0.05 with Benjamini-Hochberg correction for multiple comparisons). Large empty symbols indicate scaling and large full symbols indicate non-scaling interventions (based on the AFT+KS ^30^, see Methods, Supp. Data 1). **b**, Relative longevity and steepness of selected interventions from the literature outside of the NIA ITP, including those based on transgenic mice. Due to the varying sample size and statistical power of the studies analyzed, only interventions that are without counterparts in NIA ITP are shown. See Supp. Fig. 6 and Supp. Data 2 for all non-NIA-ITP interventions analyzed. Graphic symbols and legend are the same as those in (**a**). Round symbol denotes a study that only reported mixed sex cohorts. Effect sizes are not directly comparable between the shown interventions and those in NIA ITP because of the differences in controls. **c**-**h**, Survival curves and rescaled survival curves of steepening interventions (red) and their respective control (black). **c**, Senolytic treatment (genetic ablation of p16 expressing cells) ^22^ in mice from both sexes starting at 12 months of age. **d**, Transgenic VEGF female mice with mildly increased systemic VEGF level via induced production by hepatocytes ^59^**. e**, Acarbose (2500 ppm) in male mice starting at age of 8 months ^54^. **f**, Non-feminizing estradiol 17-α-estradiol (17a-E2) in male mice (14.4 ppm) starting from age of 16 months ^51^. **g**, Hypoglycemic and ketogenic agent 1,3-butanediol (BD) at 100,000 ppm in male mice starting at age of 6 months ^62^. **h**, angiotensin-converting enzyme inhibitor Captopril (Capt) at 180 ppm in male mice starting from age of 5 months ^62^.

Among the 65 interventions analyzed (42 in NIA ITP, 26 outside of NIA ITP) most did not show significant steepening effects - they showed approximate scaling (AFT+KS test, for p-value, see Supp. Data 1,2). We identified several classes of interventions that showed significant lifespan increase with steepening survival curves (Fig. 4a-b, Supp. Fig. 4). The steepening intervention classes include senolytic treatment, ketogenic diet and VEGF overexpression, and, in males only, 17α-estradiol ^51–53^ and hypoglycemic agents such as acarbose ^54,52,53^ and canagliflozin ^55^ (Fig. 4c-h).

In the NIA ITP dataset, acarbose ^54^ and 17α-estradiol ^51^ showed a dose dependent effect, with both lifespan and steepness increasing with dose. Additional interventions showed steepening with only mild lifespan extension, such as green tea extract (GTE) (females) ^56^ and nordihydroguaiaretic acid (NDGA) ^57^ (males) in the NIA ITP data (Fig. 4a, Supp. Fig. 7). The antioxidant taurine has a mild steepening effect (Supp. Fig. 6). High dosage of rapamycin (42ppm in females) causes steepening but is different from other rapamycin data points which show approximate scaling (Fig. 4a, Supp. Fig. 8, see Methods for statistical test of scaling).

Senolytic treatments showed life extension using a genetic approach to ablate cells that express p16 ^22,58^. Transgenic production of VEGF showed a 45% life extension and 148% increased steepness for females ^59^ (Fig. 4b, Supp. Fig. 6). Cyclic ketogenic diet in two separate studies ^60,61^ (Supp. Fig. 6, Supp. Data 2), and a ketogenic dietary supplement ^62^ showed evidence of life extension and steepening.

The steepening interventions may constitute targets for future measurements of mice sickspan.

### Combinations of steepening and scaling interventions show additive e<ects consistent with the SR model

One application of the present approach concerns combinations of interventions. Intuitively, interventions that affect different pathways, such as damage production and damage removal, should act independently on survival. The SR model helps pinpoint such interventions - for example an intervention with scaling and an intervention with steepening are predicted to affect different pathways. Combining a perturbation that scales with one that steepens should therefore provide the best of both worlds - a steep survival curve in which lifespan is extended beyond either individual intervention. This prediction is supported by NIA ITP data on a combination of acarbose (steepening in males) and rapamycin (scaling) administered at 9 months. Longevity was enhanced compared to acarbose or rapamycin alone, and steepness was comparable to the acarbose mono-treatment (Fig. 5). Simulations of the SR model calibrated to these interventions corroborate this phenomenon (Supp. Fig. 9).

**Figure 5.**
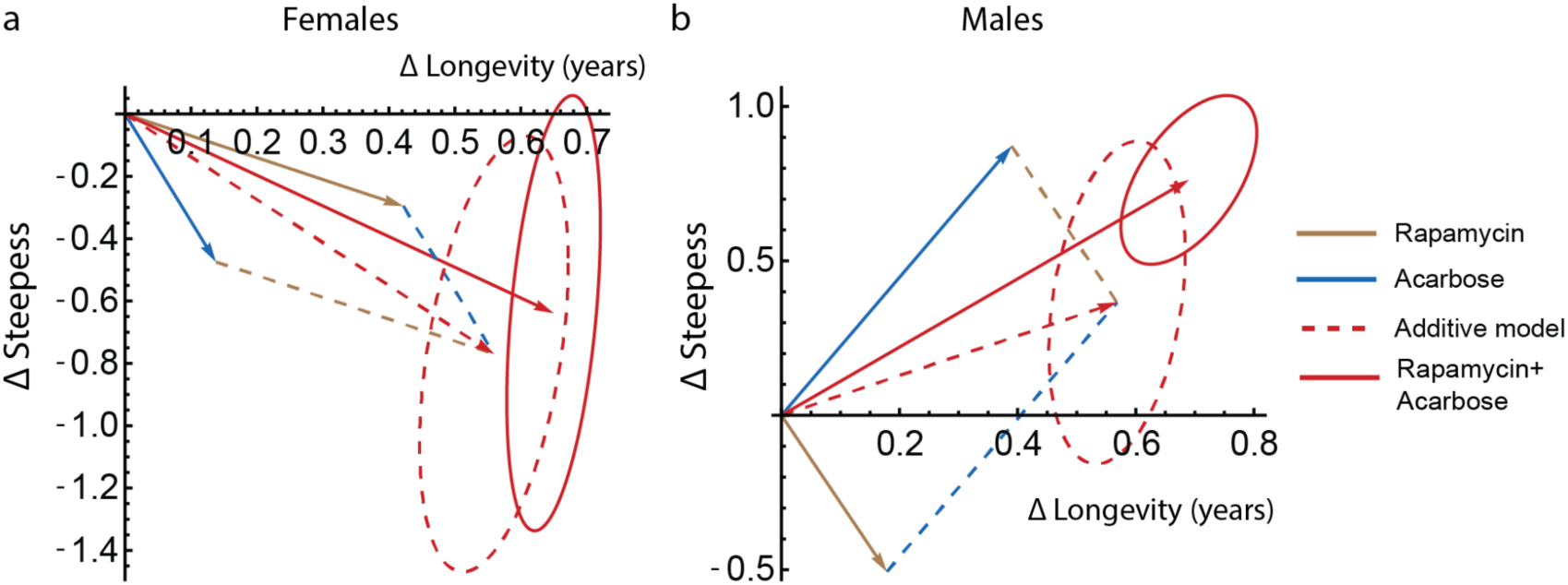
Combination data supports SR model prediction that interventions that target production and removal extend lifespan and steepen additively. Change in longevity and steepness in the ITP data for acarbose (1000 ppm, starting at 8 months of age), rapamycin (14 ppm, starting at 9 months of age) and their combination (Rap 14.7 ppm + Ac 1000 ppm starting at 9 months of age). **a**, female mice, **b**, male mice. Ellipses are 95% CI (bootstrapping). Dashed lines and ellipses are additive effects.

### Steepening of human survival curves is largely due to reduction in extrinsic mortality

The present study primarily aims to understand lab organism data. Considerations in humans are far more complex because as compared to model organisms with defined genetics living in controlled environments, sickspan is affected by additional variations in lifestyle, environment, socioeconomic status and medical care. We therefore provide a tentative account of how the present considerations might apply to human data.

There has been a global trend of rectangularization of human survival curves in the past century ^7^ -- in many countries, the median lifespan has increased and survival curve steepness has increased as well ^63^ (Fig. 6a). There is a tight correlation between life expectancy and lifespan equality (a measure of steepness) ^64^ . There are also exceptions in certain countries in which lifespan variability has increased or remained constant in recent decades ^63^.

**Figure 6.**
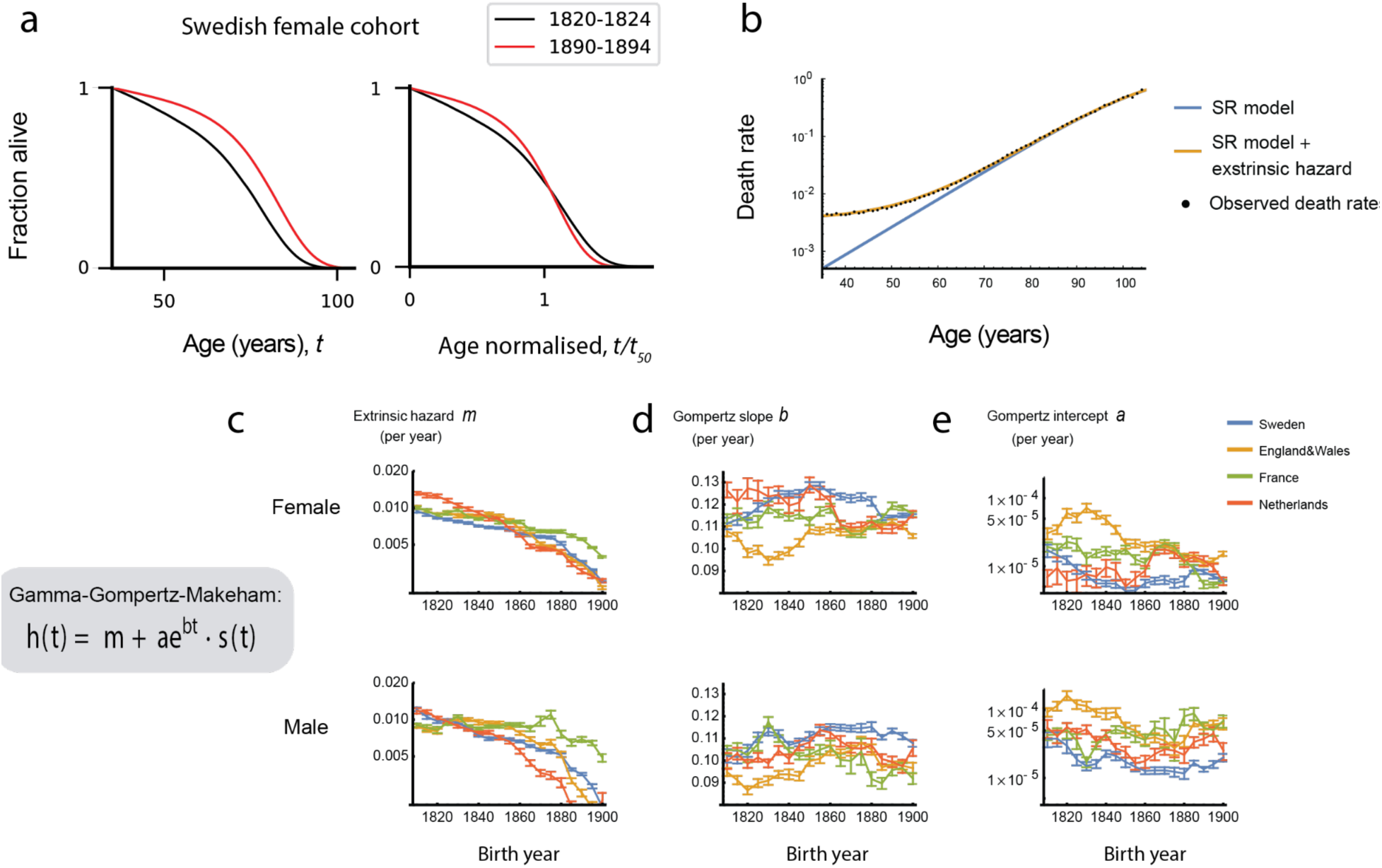
Steepening (rectangularization) of human survival curves in the past century is mainly driven by declining extrinsic mortality. **a**, Survival curves and rescaled survival curves of 2 cohorts of Swedish women born in 1820-1824 and 1890-1894. **b**, Hazard curve (death rates by age) of Swedish women born in 1890-1894, and of SR model fitted to this cohort. The SR model parameters that correspond to the blue and orange hazard curves are *η*=1.04x10^-5^/day^2^, *β*= 0.45 /day, *κ*=0.50, *ε*=0.48 /day, *Xc*=19.9 ^32^. The extrinsic hazard (Makeham term) in the orange curve is m=0.0036 per year. **c-e**, Mortality component trends of Swedish, Dutch, French (civilian) and English & Welsh combined (civilian) cohorts born 1810 to 1910. Extrinsic hazard rates, Gompertz slopes and Gompertz intercepts are estimated using the Gamma-Gompertz-Makeham model ^68^. In the SR model, Gompertz slope can be approximated by *ηXc/ε* and intercept is proportional to e*^βXc/ε^* ^32^. The Gamma- Gompertz correction term s(t) is s(t)=e^c^/(e^c^-1+e^bt^), where c is an additional parameter that ranges from 7.5 to 10, such that s(t) only comes into effect at old age.

Relative sickspan has not been as extensively measured as lifespan, with data accumulating only recently. Available data indicates that relative sickspan since 1980 has compressed in some countries and has grown in others ^8,65^.

We asked whether the rectangularization might be due to changes in SR model parameters, to test whether such changes might predict relative sickspan in each country. To do so, we analyzed human cohort mortality data from the Human Mortality Database (HMD) ^66^. Human data shows a sizable extrinsic mortality that dominates mortality until age 30-40. In order to account for this, we fitted human hazard curves to the SR model with an added extrinsic mortality term (Fig. 6b). The fits are excellent (adjusted R^2^=0.9994, F-test p<10^-16^, see Fig. 6b).

We find that the main driver for rectangularization of the survival curve in human data is the decline in extrinsic morality. Similar results are found when fitting human hazard to the Gamma-Gompertz- Makham equation ^67,68^ -- the age independent extrinsic mortality drops with cohort birth year (Fig. 6c), whereas the Gompertz slope and intercept remain nearly constant (Fig. 6d,e). The dominance of the extrinsic mortality drop in human cohorts is consistent with previous studies ^7,9^. This contrasts with model organism data analyzed above in Fig. 3. Unlike human populations, these animal cohorts were studied in controlled environments, and extrinsic mortality is not the main determinant of their survival curve steepness.

In addition to changes in extrinsic mortality, there have been age-specific improvements in mortality rates. Until recent decades this was primarily a reduction in young deaths (younger than median lifespan), and in the past few decades this shifted to a reduction in old deaths (older than median lifespan) ^69^. These changes are likely to be due to public health and medical interventions, as well as changes in education, socioeconomic status and other factors ^70^. Such age-specific changes confound an attempt to fit changes in SR model parameters to this human data. Thus, why some countries show a reduction in relative sickspan whereas others show an increase ^65^ may not be directly linked to changes in SR model parameters.

## Discussion

We present the hypothesis that longevity interventions that steepen survival curves compress relative sickspan. In contrast, interventions that scale the survival curve extend healthspan and sickspan proportionally as they extend lifespan, and thus do not compress relative sickspan. These effects hold in the saturating removal mathematical model of aging, where interventions that affect damage production cause approximate scaling and mild extension of sickspan, whereas interventions that increase damage removal cause steepening and compress sickspan. The hypothesis is validated in data from invertebrates and mice. We discuss interventions in mice which steepen the survival curve as candidates to compress relative sickspan. The theory also applies to combinations of interventions showing additive effects in interventions that affect steepness and lifespan.

One can intuitively argue for this hypothesis without need for a mathematical model. Many of the longevity interventions that scale the survival curve seem to generally slow the tempo of aging processes (e.g. temperature in invertebrates, dietary restriction). If one assumes that age-related morbidity originates from the same factors that drive aging and mortality, it makes sense that healthspan and sickspan should also scale by the same proportions as lifespan.

In contrast, longevity interventions that steepen the survival curve may be argued to affect processes more specific to aging-related decline than its overall tempo. They make the mortality curve more rectangular and thus the time of death less variable between individuals. This can be considered to reduce the proportion of frail individuals that normally die at ages younger than the median lifespan. Steepening interventions seem to make the whole population more robust (longer lived) and at the same time more homogenous in terms of frailty. An extreme steepening means that almost all individuals reach the maximal lifespan. In humans, individuals close to the maximal lifespan often have a relatively short sickspan ^71,72^. Thus, it is plausible that relative sickspan is shortened by steepening interventions.

To obtain a quantitative account of how steepness may result in compressed sickspan we used the saturating removal (SR) model. This model was developed and calibrated based on data in mice and adjusted to human timescales^32^. The SR model assumes a form of damage *X* that drives morbidity and mortality, which rises with age in a fluctuating way. The model is agnostic to the nature of this damage, which could be different in different species. The model includes production of driving damage *X* at a rate that rises linearly with age - corresponding to accumulation of damage-producing units that are not removed. It also assumes that removal of the damage *X* occurs by mechanisms that saturate at high damage.

The SR model was previously validated against patterns of aging from diverse studies. The model displays the Gompertz law of mortality in which hazard rises exponentially with age, as well as the late age slowdown of hazard, and explains the Weibull-like shape of mice survival curves ^32^. The model quantitatively predicts the effect of mice parabiosis on senescent cell parameters ^33^ in agreement with data. Parabiosis in the model is explained by the young mouse sharing removal capacity with the old mouse (e.g. immune cells). The SR model predicts the slowdown in senescent cell removal rate with age in agreement with mice experiments ^32^. It also explains the rapid shifts between survival curves upon dietary midlife interventions in flies, and the change in hazard slope upon temperature changes ^32^. The SR model was extended by Katzir et al to explain in quantitative detail the exponential rise of human age-related disease incidence, as well as the incidence decline at very old ages. The extended model assumed two parameters for each disease - a disease threshold and a fraction of susceptible individuals - and provides excellent fits to the incidence curve of hundreds of age-related diseases from large medical record datasets ^34^.

Here we show that the SR model predicts that perturbations that increase the steepness of survival curves also compresses the sickspan, whereas any perturbation that scales the survival curve also approximately scales the sickspan (extending relative sickspan mildly). More specifically, changing the rate of production of the damage that drives aging leads to approximate scaling of mortality and sickspan. Changing other parameters, such as raising the specific removal rate of the driving damage, extends lifespan with steepening and compresses morbidity.

One exception is raising the death threshold without correspondingly raising the disease threshold. This steepens the survival curve, but increases relative sickspan, since individuals spend more time between the disease and death thresholds. As a thought experiment, consider artificial life expansion by intensive care units - such perturbations increase sickspan by delaying death. A steep survival curve in these situations can result by artificially keeping individuals alive to the same age. We assume that the longevity interventions in the controlled experiments in model organisms that we analyzed here do not cause such uncoordinated changes in death and disease thresholds. In humans, sex differences in the death and morbidity thresholds between females and males could explain the’ health survival paradox’ - that females have longer lifespans but expanded morbidity. This would occur if *Xc* is higher in females (and *Xd* is either the same or lower in females).

The SR model makes a major simplifying assumption—that a single damage variable *X* determines both morbidity and mortality. This provides quantitative predictions that can be tested against data, as in Fig. 3,6. Future work can explore more complex situations — for example that damage in each organ affects health differently from lifespan. In such a case there might be multiple qualitatively distinct damage variables rather than a single *X*. There is evidence that health and lifespan may have overlapping but distinct determinants, and that cognitive functions may have somewhat different aging patterns than other physiological functions ^34,41,73–75^ . Furthermore, on the population level, genetic and environmental heterogeneity can affect the onset of morbidity and complicate interpretation of the model. Extensions of the SR models can in principle address this complexity ^34^.

### Survival curve shape as a tool to interpret the mechanism of longevity interventions

Aging research has made progress in understanding molecular and cellular processes that contribute to aging. There is a need for theoretical frameworks to integrate these empirical findings ^76^. The present framework offers a tool to interpret longevity interventions using the shape of the survival curve: Scaling points to effects on production of the driving damage, whereas steepening or shallowing points to effects on removal of this specific damage or its death threshold. We next discuss how this approach points to specific factors as candidates for the driving damage X.

Senolytics extend lifespan and steepen the survival curve, as noted by Kowald and Kirkwood ^58^ who analyzed data from two types of senolytic drugs and from genetic ablation of p16 positive cells. In our analysis, their effect resembles increasing removal rate *β* in the SR model. Thus, in mice, senescent cells (and more generally damaged cells) may play the role of the driving damage *X* in the SR model, as suggested by Karin et al ^32^.

An intriguing class of steepening interventions includes cyclic ketogenic diet ^60,61^, the ketogenic agent 1,3-butanediol (BD) and the diabetes drugs acarbose ^54^ and canagliflozin ^55^. These interventions share a common mechanism: they all lower glucose spikes, the fraction of time where glucose is at high levels. The drug acarbose inhibits an enzyme that releases glucose from complex carbohydrates in the gut. canagliflozin is an SGLT2 inhibitor that inhibits kidney glucose reabsorption. These interventions extend median lifespan in male but not female mice.

Glucose spikes are thought to primarily damage the vasculature. Damage to blood vessels is a major pathology in diabetes, causing cardiovascular and renal disease. Endothelial cells are damaged by high glucose in several ways including reactive oxygen species, glycation of proteins and activation of signaling pathways ^77^. Intermittent high glucose causes senescence in endothelial cells ^78^ .

The vasculature-protecting mechanisms appear in our analysis to increase damage removal rate *β*. One explanation is that damaged microvasculature reduces access of immune cells to the senescent and damaged cells in tissues. Interventions that protect microvasculature may thus enhance the ‘roads’ for immune clearance, and hence increase the removal rate *β*. This can contribute to their steepening effect on the survival curves.

The notion of vascular protection as a mechanism for the steepening interventions is consistent with a VEGF longevity intervention which increased mouse lifespan by 40% and, in females, steepens the survival curve ^59^. VEGF promotes vascular repair and slows the age-related loss of microvasculature. Vasculature protection may also be relevant for captopril (a hypertension drug), which increases steepness with a mild longevity gain.

The non-feminizing estradiol, 17-α-estradiol, steepens survival curves and extends lifespan even when administered late in life in male mice ^79^; it has no longevity effect in females. The longevity effects of this estradiol are reported to depend on testicular hormones. It is not clear how this estradiol ties into the mechanism of the other steepening interventions, although it is plausible that it has protective roles on tissues ^80,81^.

Rapamycin treatment also extends life, even when started at middle age ^53,82–85^, but does not generally steepen the survival curve except at the old-age tail of the survival curve (Supp. Fig. 8). It may thus have a primary effect that scales survival, such as mTOR inhibition. Notably, one experiment with rapamycin at high dose shows a steepening effect in females, particularly on the part of the survival curve that corresponds to very old ages. It may thus have a secondary effect at high dosage on older individuals.

Additional interventions that cause steepening but with milder longevity effects include antioxidants and anti-inflammatory agents (NDGA ^57^, green tea extract ^56^) (Supp. Fig. 7). In the SR model, strong steepening with mild lifespan extension characterizes coordinated increases in the mortality and morbidity threshold *Xc* and *Xd*. One may hypothesize that these thresholds are affected by tolerance to inflammatory damage which is a major causal factor in aging ^86,87^. Indeed, a major deleterious effect of senescent and damaged cells is secretion of inflammatory factors ^88^.

Interventions that scale the survival curve are predicted not to compress sickspan - instead they are suggested to extend sickspan by the same proportion that they extend lifespan. These interventions include caloric restriction. Caloric restriction has pleiotropic effects beyond its effects on longevity such as reducing body weight, reducing fertility and enhancing frailty ^89^.

It would be interesting to see whether emerging interventions such as in vivo partial epigenetic reprogramming scale or not. Data from preliminary studies does not show significant steepening ^23,24^ (Supp. Fig. 10), although further research is warranted.

Future work can further explore how the SR model relates to human mortality and sickspan. Rectangularization of the human survival curve over the past century is primarily due to reduction in extrinsic mortality rather than changes in intrinsic parameters. Old-age survival and morbidity in humans is impacted by heterogeneity of socioeconomic status ^90–92^, lifestyle, genetics ^72^, medical care and public health ^92^, as well as emotional well-being - older cohorts appear to have high average well-being ^93^, whereas social isolation in the old has major negative effects ^94^. Such factors can complicate a direct comparison to the SR model.

## Summary

We present a theory and evidence that longevity interventions that steepen the survival curve compress morbidity. Conversely, interventions that do not change the shape of the survival curve, namely interventions that show scaling, extend both lifespan and healthspan proportionally and do not compress morbidity. The theory is supported by data from model organisms and by mathematical modeling using the saturating removal (SR) model of aging. The theory helps to interpret intervention mechanisms of action by means of survival curve shape - we discuss various interventions in mice that steepen survival curves, such as senolytics, ketogenic diet, acarbose, and canagliflozin, and suggest their potential mechanisms of action. We apply this to combinations of interventions. This study provides insights into the relationship between survival curves, longevity interventions, and relative sickspan, contributing to our understanding of strategies to improve health in aging individuals.

## Methods

### Saturating removal (SR) model simulations

We simulated the SR model using parameters reported in Karin et al ^32^. For mice, *η*=2.3x10^-4^/day^2^, *β*=0.15 /day, *κ*=0.5, *ε*=0.16 /day, *Xc*=17. For human, the baseline parameters are *η*=1.4x10^-3^/(day*year), *β*=0.15 /day, *κ*=0.5, *ε*=0.16 /day, *Xc*=17. The sickspan threshold *Xd* was set to 12, which is the lower end of the estimated thresholds for age-related diseases ^34^. This results in a mean sickspan of 12.3y, consistent with reported age achieved without major chronic diseases in the United States ^95^. We simulated interventions by varying an individual parameter *η*, *β*, *ε or Xc,* or by simultaneously changing *Xc* and *Xd* by the same factor. Simulation used Mathematica 14.0.

### Survival and healthspan data

Individual-based longitudinal survival and health data are collected from references ^40–42^ . The authors of reference ^40^ provided the raw data. Data from reference ^41^ and reference ^42^ was downloaded from github https://github.com/nstroustrup/2022_hierarchical_process_model and figshare (DOI: 10.6084/m9.figshare.25125587) respectively. Survival and healthspan distributions from references ^26^ were obtained by digitizing relevant figures using WebPlotDigitizer (https://apps.automeris.io/wpd/). In Fig. 3d one outlier datapoint was removed: glp-1 in Oswal et al (relative steepness 1.08±0.11, relative sickspan 0.48±0.05, Supp. Table 4). This mutant was also tested in Statzer et al and included in Fig. 3c, where a different sterile strain was used as control.

For experiments conducted by NIA ITP, we downloaded lifespans of individual mice from all experimental and control cohorts (C2004 - C2017) directly from the Mouse Phenome Database (https://phenome.jax.org/projects/ITP1, June 2023). Experiments with different drug concentrations and different initiation age and regimes of exposure are analyzed and reported separately as different interventions. Data from all 3 sites of NIA ITP were pooled together. NIA ITP used UM-HET3 mice, a heterogenous stock produced by a four-way cross.

For mice experiments independent of NIA ITP, we downloaded the PDFs of the relevant publications and used WebPlotDigitizer (https://apps.automeris.io/wpd/) to digitize the survival curves. We also recorded the genetic background and the sample sizes of control and treated cohorts. From these survival curves, we inferred the time of deaths of individual mice constituting the data. We excluded publications using mice strains considered to be disease models that have specific deleterious genes. In contrast to NIA ITP, all but one study used inbred strains or hybrids involving 2-way or 3-way crosses among inbred strains.

Human hazard curves are downloaded from the Human Mortality Database ^66^. Relevant health expectancy surveys are described in reference ^8,65^ and digitized from references ^96,97^.

### Statistical analysis of mice survival curves

We calculated steepness from the survival curves defined as the median lifespan divided by the interquartile range (IQR), that is t50/(t75-t25), where tx the age at which x% survive. This definition of steepness is the inverse of relative IQR studied by Carnes et al ^29^.

To ascertain whether an intervention changes the shape of the survival curve, we follow Stroustroup et al ^30^ and use an accelerated failure time (AFT) approach with a Kolgomorov-Smirnov (KS) test modified to accommodate right censored data ^98^. The rescaling is based on the accelerated failure time model implemented using Python package lifelines (version 0.27.7). We use this test to classify treatments into scaling and non-scaling categories. We apply a threshold on the KS effect size of 0.05 (Fig. S4).

For Fig. 4ab, we sought to identify interventions with a significant effect in terms of t50 and steepness. For this purpose we bootstrapped the death times with returns. For every mice cohort (control and interventions per ITP site and sex), we resampled with returns the recorded death times and censorship status. We repeated this 200 times. Based on these bootstrapped samples, we estimated the joint distributions of median lifespan and steepness for both control *μc*=(ln(*Lc*), ln(*Sc*)) and intervention cohorts *μi*=(ln(*Li*), ln(*Si*)) by 2D normal distributions *Nc*(*μc*, *σc*) and *Ni*(*μi*, *σi*). We then tested the null hypothesis that the control and intervention groups have the same longevity and steepness (H0: *Li*/*Lc*=1 and *Si*/*Sc*=1) using Hotelling’s t-squared statistic. Because *μi* - *μc* ∼ *Ni*(*μi*-*μc*, *σi*+*σc*), H0 is equivalent to a Mahalanobis distance between control group (ln(*Lc*), ln(*Sc*)) and intervention group (ln(*Li*), ln(*Si*)) of 0. Thus, for every intervention and its control, we calculated the p-value that *μi* - *μc*=0 using Hostelling’s t-squared statistic and Hotelling’s T-squared distribution with 2 dimensions and 198 degrees of freedom.

For intervention experiments conducted by NIA ITP, we apply a p-value threshold adjusted for multiple comparisons using the Benjamini-Hochberg method. For an α level of 0.05, the adjusted p-value threshold is 0.0113. For intervention experiments conducted independent of NIA ITP, due to the varying data quality and statistical power, we conducted Benjamini-Hochberg adjustment separately from those in NIA ITP, resulting in an adjusted p-value threshold of 0.0214 (α=0.05).

### Statistical analysis of human mortality data

We downloaded human cohort mortality data from the Human Mortality Database ^66^. We used the cohort death rate and exposure-to-risk data with 1x5 (age x year) intervals to estimate model parameters.

We estimated the extrinsic (Makeham) and age-related (Gamma-Gompertz) components of mortality for each cohort using the Gamma-Gompertz-Makeham model h(t)=m+a*e^c+bt^/(e^c^-1+e^bt^). We fitted the log-transformed age-specific death rates using the ‘NonlinearModelFit’ of Mathematica 14.0. We used the square root of death numbers as weights for each age group, to balance the strengths of evidence with residue minimization in the log hazard space. The goodness-of-fit are excellent for the countries and cohorts shown in Fig.6c-e, with adjusted-R^2^ > 0.995.

We identified SR model parameters that correspond to a given cohort by minimizing the residues in log hazard space, between Gamma-Gompertz hazards h(t)=a*e^c+bt^/(e^c^-1+e^bt^) of a given cohort, and hazards of simulated SR models. Like the fitting metric above, the residues were weighted according to the square root of death numbers. The simulation was done in Mathematica 14.0, using compiled code implementing the Euler-Maruyama method. Each simulation was done with N=5*10^4^ simulated individuals.

