## Supplementary Figures 1-10, Supplementary Tables 1-4, Supplementary Data 1-2 for "Compression of morbidity by interventions that steepen the survival curve"

#### Table of contents

S1. Effect of SR model parameter variation on longevity and sickspan 2
S2. Tables of mice, flies and worms control and intervention cohorts with longitudinal healthspan data 3
S3. Effect of steepness on mice sickspan is robust to the definition of sickspan threshold 6
S4. Several interventions in the NIA database significantly change the steepness of survival curves 7
S5. The prevalence of steepening interventions for male mice and their shorter lifespan and shallower survival curves 9
S6. Intervention experiments independent of NIA ITP introduce new classes of steepening interventions and agree with those in the NIA ITP database 10
S7. Antioxidants NDGA and GTE increase steepness 12
S8. Rapamycin shows approximate scaling except at old ages 13
S9. Rapamycin and acarbose interact additively to extend mouse lifespan 14
S10. In vivo reprogramming shows approximate scaling in preliminary experimental data 15

### S1. Effect of SR model parameter variation on longevity and sickspan

Here instead of lifespan on the axis axis as in Fig. 1e, Fig. 2 cd, we plot the log magnitude of parameter change on the x axis. Baseline parameters are for mice [32] .

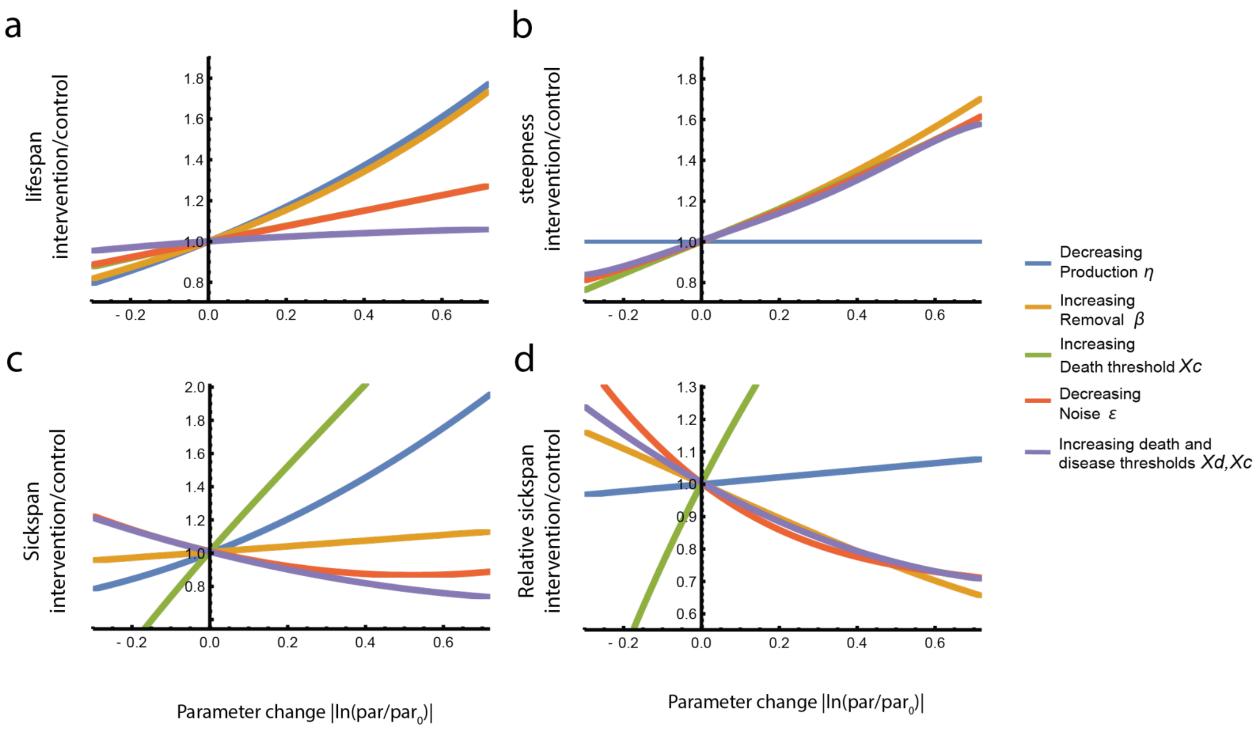


**Supplementary Figure 1. Effect of SR model parameter variation on longevity, steepness and sickspan.** Lifespan, steepness, absolute sickspan and relative sickspan are plotted against magnitude of parameter changes in the direction indicated by the legend. Simulations are the same as in Fig. 2b-d. Parameters are shown relative to the baseline parameter set (see Methods, $\eta$=6.6×10$^{-6}$/day$^2$, $\beta$=0.15 /day, $\kappa$=0.5, $\varepsilon$=.16 /day, $Xd$=12, $Xc$=17). The same qualitative effects are seen for other parameter choices.

### S2. Tables of mice, flies and worms' control and intervention cohorts with longitudinal healthspan data

In all cases (Supp. Table 1-4), sickspan is defined according to the original paper criteria. The criteria for each paper is briefly summarized in table captions.

| strain | sex | diet | longevity (weeks) | longevity S.E. (weeks) | steepness | steepness S.E. | sickspan (weeks) | sickspan S.E. (weeks) | relative sickspan | relative sickspan S.E. |
|---|---|---|---|---|---|---|---|---|---|---|
| C57BL/6J | f | AL | 127.9 | 3.9 | 9.2 | 3.5 | 2.5 | 0.7 | 1.9% | 0.5% |
| C57BL/6J | m | AL | 131.1 | 6.2 | 4.8 | 0.9 | 9.6 | 1.7 | 6.8% | 1.1% |
| J:DO | f | AL | 149.3 | 2.8 | 11.5 | 3.3 | 7.5 | 1.4 | 4.8% | 0.8% |
| J:DO | f | CR20 | 168.5 | 6.0 | 5.2 | 0.6 | 12.3 | 1.4 | 6.9% | 0.7% |
| J:DO | f | CR40 | 178.4 | 5.4 | 4.4 | 0.5 | 19.9 | 1.7 | 10.4% | 0.8% |
| J:DO | f | IF1 | 151.0 | 2.7 | 10.1 | 2.1 | 6.6 | 0.9 | 4.2% | 0.6% |
| J:DO | f | IF2 | 157.2 | 4.2 | 6.7 | 0.9 | 6.4 | 0.8 | 4.0% | 0.5% |

**Supplementary Table 1. Longevity, steepness and sickspans of mice cohorts under dietary longevity interventions**
**(caloric restriction and intermittent fasting) in two strains from Luciano et al [42].** Sickspan was defined using mouse frailty index. In this table, female J:DO is considered to be the control cohort. Comparing steepness and relative sickspan of all other cohorts to those of female J:DO generates the intervention/control steepness and relative sickspan shown in Fig. 3a.

| Group | Sex | N survival | N sickspan | longevity (days) | longevity S.E. (days) | steepness | steepness S.E. | sickspan (days) | sickspan S.E. |
|---|---|---|---|---|---|---|---|---|---|
| Oregon (control) | female | 115 | 105 | 64.2 | 0.6 | 8.8 | 0.9 | 0.79 | 0.12 |
| Lausanne | female | 93 | 83 | 33.0 | 1.0 | 2.9 | 0.6 | 1.68 | 0.20 |
| Protein restriction (PR) | female | 122 | 112 | 84.9 | 1.5 | 5.3 | 0.7 | 0.86 | 0.11 |
| Curcumin | female | 95 | 85 | 66.5 | 1.6 | 4.6 | 0.6 | 1.68 | 0.21 |
| Superfruit | female | 88 | 79 | 69.4 | 2.0 | 4.0 | 0.4 | 1.33 | 0.24 |
| PR+Cur | female | 94 | 82 | 76.4 | 1.1 | 7.0 | 1.3 | 1.59 | 0.23 |
| PR+SF | female | 119 | 109 | 76.2 | 1.1 | 5.2 | 0.5 | 1.50 | 0.23 |

| | | | | | | | | | |
|---|---|---|---|---|---|---|---|---|---|
| Oregon (control) | male | 109 | 94 | 71.2 | 2.2 | 4.0 | 0.4 | 1.12 | 0.19 |
| Lausanne | male | 98 | 90 | 51.2 | 2.0 | 3.0 | 0.3 | 1.43 | 0.15 |
| Protein restriction (PR) | male | 113 | 102 | 87.2 | 1.4 | 5.1 | 0.3 | 0.58 | 0.10 |
| Curcumin | male | 87 | 79 | 84.5 | 2.0 | 4.7 | 1.0 | 1.68 | 0.24 |
| Superfruit | male | 110 | 101 | 85.9 | 2.1 | 4.0 | 0.7 | 1.06 | 0.15 |
| PR+Cur | male | 104 | 97 | 81.9 | 2.0 | 4.9 | 0.5 | 0.84 | 0.13 |
| PR+SF | male | 99 | 87 | 87.1 | 1.5 | 6.1 | 0.8 | 0.57 | 0.09 |

**Supplementary Table 2. Longevity, steepness and sickspans of *D. melanogaster* cohorts under nutritional longevity interventions in two strains (Lausanne and Oregon)**. Sickspan was defined by deficits in climbing, jumping and flying in response to a mechanical challenge. Fig. 3b is generated using Oregon male and female cohort controls for all other male and female cohorts respectively. This table is generated from raw data provided by the original authors from Gaitanidis et al [40].

| Group | Longevity (days) | Steepness intervention/ control | Steepness intervention/ control C.V. | Sickspan (days) | Sickspan lower C.I. (days) | Sickspan upper C.I. (days) | relative sickspan intervention/ control | relative sickspan intervention/ control lower C.I. | relative sickspan intervention/ control upper C.I. |
|---|---|---|---|---|---|---|---|---|---|
| N2 (wild type) | 31.2 | 100.0% | 10.2% | 21.4 | 10.7 | 24.7 | 100.0% | 49.8% | 115.6% |
| daf2(e1368) | 42.6 | 189.6% | 9.3% | 15.4 | 10.7 | 18.4 | 52.6% | 36.5% | 62.9% |
| daf2(e1370) | 44.4 | 171.7% | 12.7% | 24.7 | 15.8 | 27.0 | 81.0% | 51.8% | 88.7% |
| eat2(ad1116) | 31.9 | 113.5% | 10.0% | 14.2 | 9.7 | 18.9 | 64.7% | 44.5% | 86.3% |
| glp1(e2141) | 34.7 | 212.2% | 9.2% | 7.9 | 6.1 | 8.8 | 33.3% | 25.6% | 36.9% |

**Supplementary Table 3. Longevity, steepness and sickspan of longevity mutants in *C. elegans* where sickspan was defined by loss of muscle power measured by piezoelectric challenge.** Data digitized from Statzer et al [26].

| experiment | Intervention type | Intervention description | Control group genotype | Control group food | Control group temperature | Steepness intervention/ control | Steepness intervention/ control C.V. | relative sickspan intervention/ control | relative sickspan intervention/ control C.V. |
|---|---|---|---|---|---|---|---|---|---|
| oswal_daf2 | Genotype | daf2(e1368) | QZ0(WT) | live OP50 | 25 | 140.4% | 7.4% | 85.8% | 3.2% |
| oswal_eat2_1 | Food | Bacterial Deprivation | QZ0(WT) | live OP50 | 20 | 151.2% | 8.0% | 96.3% | 3.1% |
| oswal_eat2_1 | Genotype | eat2(ad1116 ) | QZ0(WT) | live OP50 | 20 | 159.7% | 6.5% | 102.8% | 3.0% |
| oswal_eat2_2 | Food | UV NEC937 | QZ0(WT) | live OP50 | 20 | 77.1% | 6.7% | 158.8% | 2.9% |
| oswal_eat2_2 | Genotype | eat2(ad1116 ) | QZ0(WT) | live OP50 | 20 | 69.5% | 6.0% | 129.5% | 2.9% |
| oswal_glp1 | Genotype | glp1(e2141) | QZ0(WT) | live OP50 | 20 | 108.2% | 11.4% | 47.8% | 5.4% |
| oswal_nuo6 | Genotype | nuo6(qm200 ) | QZ0(WT) | UV NEC937 | 20 | 131.5% | 17.8% | 100.7% | 5.5% |
| oswal_temp | Temperatur e | 25C | QZ0(WT) | UV NEC937 | 20 | 97.3% | 5.5% | 141.9% | 1.6% |
| statzer_sterile | Genotype | glp1(e2141) | spe9(hc88); rrf3(b26) | L4440 | 20 | 124.7% | 7.0% | 82.7% | 4.1% |
| statzer_main | Genotype | daf2(e1368) | N2(WT) | heat-killed OP50 | 15 | 187.2% | 8.9% | 114.2% | 4.5% |
| statzer_main | Genotype | daf2(e1370) | N2(WT) | heat-killed OP50 | 15 | 170.1% | 12.5% | 104.7% | 4.5% |
| statzer_main | Genotype | eat2(ad1116 ) | N2(WT) | heat-killed OP50 | 15 | 112.7% | 9.5% | 93.1% | 5.8% |
| statzer_main | Genotype | glp1(e2141) | N2(WT) | heat-killed OP50 | 15 | 211.7% | 8.8% | 80.0% | 4.6% |

**Supplementary Table 4. Genetic, dietary and environmental longevity interventions in *C. elegans*, where sickspan was measured by secession of spontaneous movements.** Data from Stazer et al [26] and Oswal et al [41].

### S3. Effect of steepness on mice sickspan is robust to the definition of sickspan threshold

In Fig. 3 we used the original author's admittedly ad hoc definition of sickspan as the time when an individual mouse exceeded 3 health deficits. Here we test the effect of changing this threshold to exceeding 4 or 5 deficits. Supp. Fig. 2a shows the raw data and these three thresholds (dashed lines at 3.5,4.5,5.5 since data is discrete).
We find that for all thresholds, relative sickspan drops with steepness of the survival curve, as in the main text (Supp. Fig. 2b).

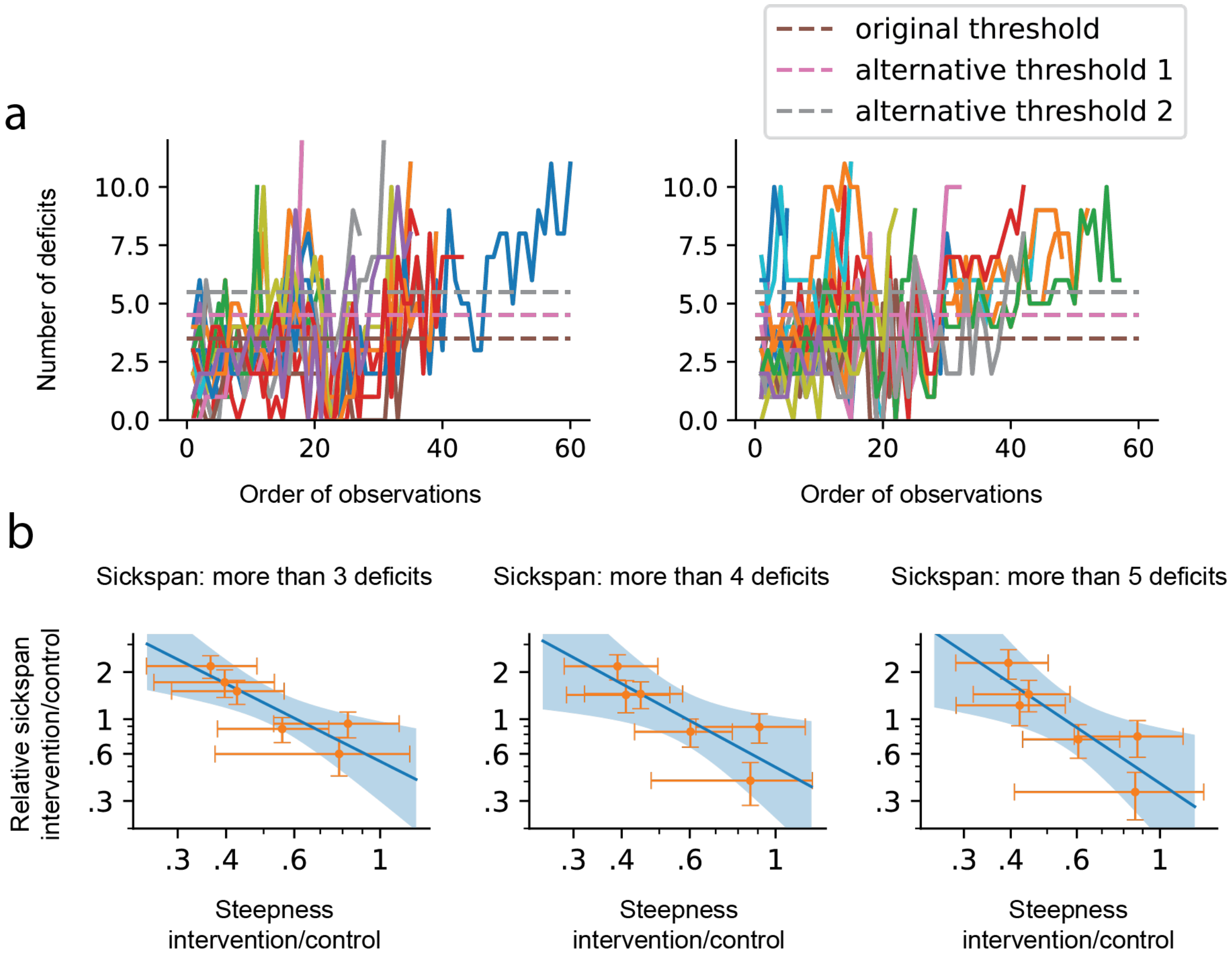


**Supplementary Figure 2. Relative sickspan in Luciano et al drops with steepness of survival curve for three different sickspan definitions, namely thresholds of above 3,4 or 5 deficits. a**, typical trajectories of mice from Luciano et al [42], C57BL/6J (left) and J:DO (right). Dashed lines indicate 3.5, 4.5, and 5.5 thresholds, since the number of deficits is integer, this corresponds to exceeding 3, 4 or 5 deficits. **b**, Analysis of relative sickspan versus survival curve steepness as in Fig. 3, for the thresholds of 3, 4, 5 deficits.

**S4. Several interventions in the NIA database significantly change the steepness of survival curves**

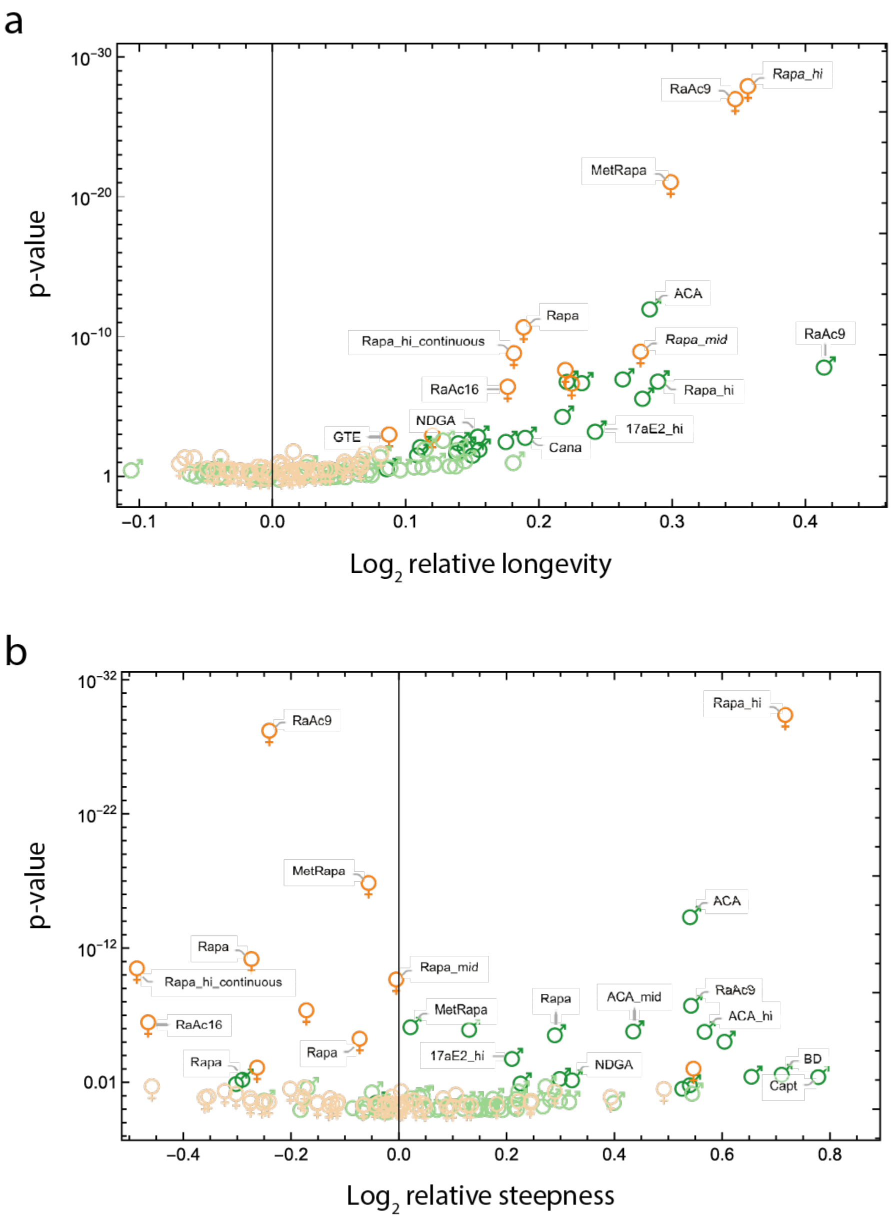


**Supplementary Figure 3. Volcano plots of all interventions in the NIA ITP database.** On y-axis, p-values of the chi-square test described in Methods (H0: treated and control groups have the same steepness and longevity) are plotted separately against effect sizes of (**a**) longevity and (**b**) steepness. All calculations are the same as in Figure 4a. Green and orange symbols indicate females and males respectively. Solid and faint symbols indicate, respectively, significant and insignificant interventions for α=0.05 with Benjamini-Hochberg corrections.

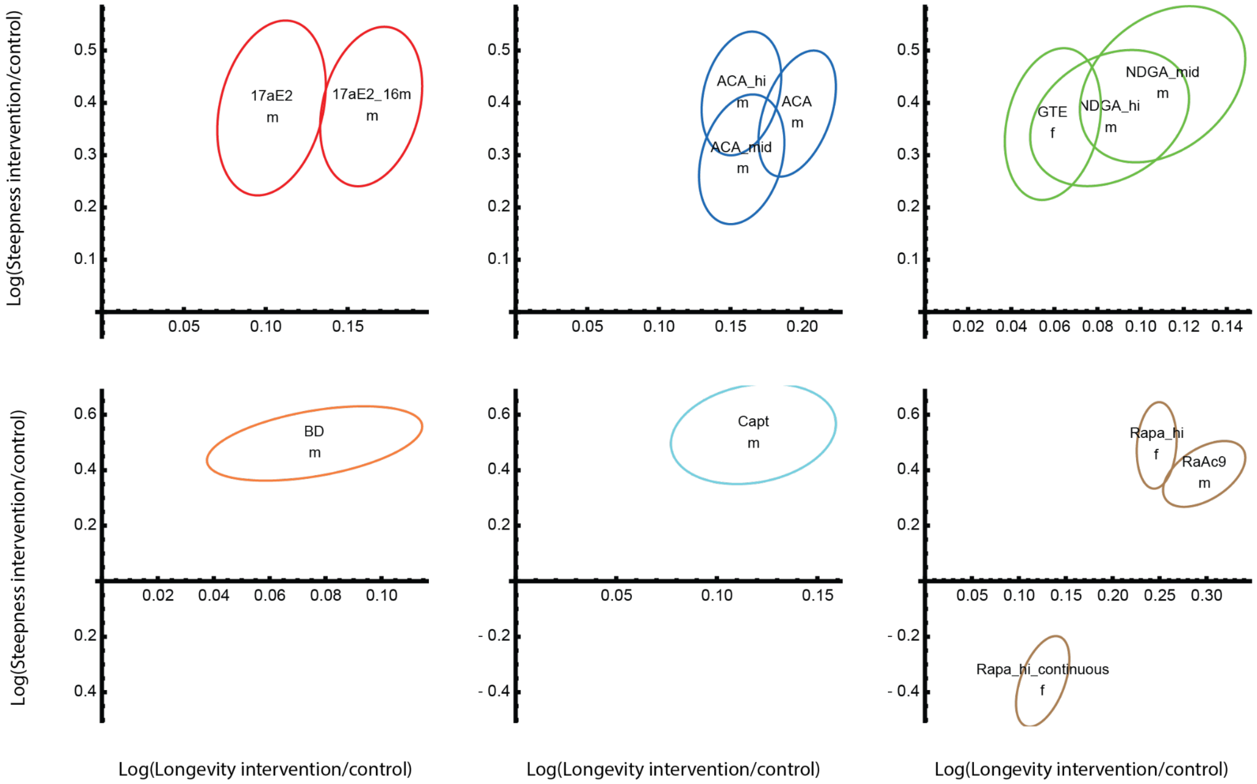


**Supplementary Figure 4. Confidence Intervals of non-scaling interventions in the NIA ITP database.** The ellipsoids indicate means and covariances of interventions' effect on longevity and steepness. For each pair of treated and cohort cohorts, the means and covariances of log(longevity) and log(steepness) of control $\mu_c$=(ln($L_c$), ln($S_c$)) and intervention cohort $\mu_i$=(ln($L_i$), ln($S_i$)) are estimated by bootstrapping 200 times with returns the right-censored death times. The mean and covariance of an intervention, $\mu_i$ - $\mu_c$ can then be modeled by multivariate normal distribution $N_i(\mu_i\text{-}\mu_c, \sigma_i\text{+}\sigma_c)$, visualized as the ellipsoids. Shown are intervention experiments found to have significant effects (Hotelling's T-square test) and non-scaling from AFT+KS test (Methods), grouped by author-assigned mechanisms of action (Fig. 4a).

**Supplementary Data 1. Statistics of all interventions in the NIA ITP shown in Fig. 4a and Supp. Fig. 3.** Relative longevity, relative steepness, quartile test p-value and AFT+KS test statistic and p-value of all interventions in the NIA ITP.

### S5. The prevalence of steepening interventions for male mice and their shorter lifespan and shallower survival curves

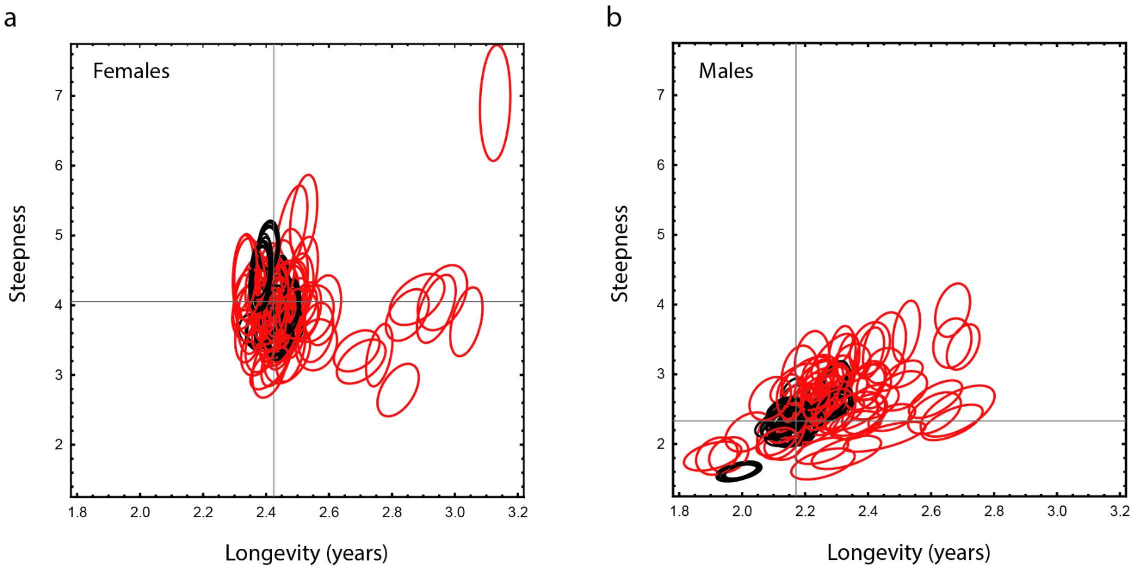


**Supplementary Figure 5. Absolute longevity and steepness of all interventions and their control groups in the NIA ITP database.** Red and black ellipsoids indicate means and covariances of treated and control cohorts respectively. For each cohort (control or treated), the means and covariances were estimated by bootstrapping 200 times the death times with returns. The crosshair cursors in each frame indicate the mean longevity and steepness of all female and male control cohorts. The ellipsoids are drawn without labeling the interventions in order to visualize the absolute longevity and steepness of all the female (**a**) and male (**b**) cohorts in the dataset.

### S6. Intervention experiments independent of NIA ITP introduce new classes of steepening interventions and agree with those in the NIA ITP database

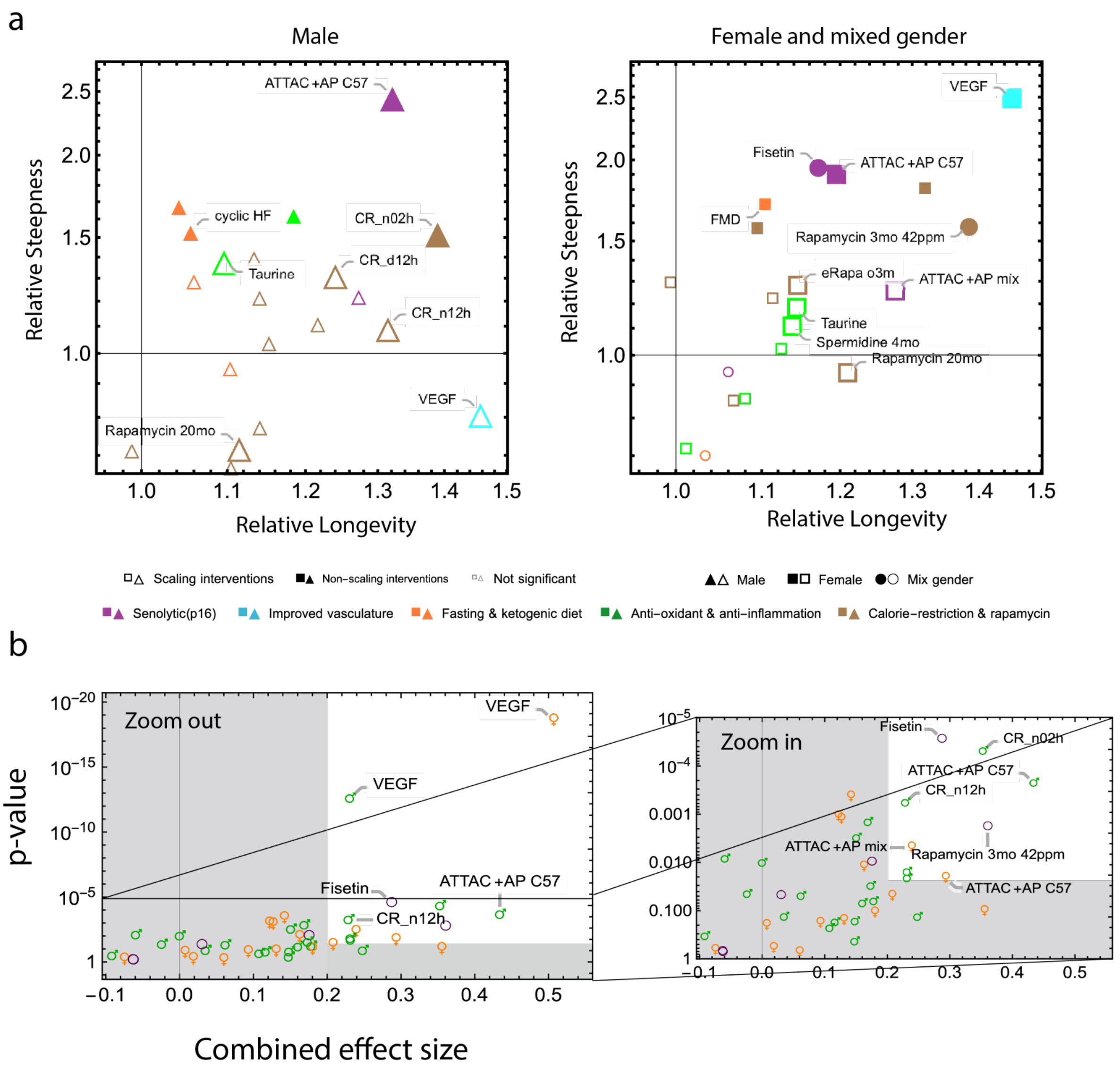

**Supplementary Figure 6. Longevity and steepness effects of Intervention not reported in NIA ITP. a**, Longevity and steepness relative to control in 44 interventions (males and females) reported in 32 publications independent of the NIA ITP (Supp. Data 2). Similar to Fig. 4a-b, triangle, square and circle symbols denote interventions for male, female and mixed gender mice cohorts respectively. Graphically styles are also similar to those in Fig. 4a: small symbols indicate interventions that are not statistically significant (quartile test, α=0.05 with Benjamini-Hochberg correction). Filled and empty symbols indicate scaling and non-scaling interventions respectively (based on a relative steepness threshold of 1.45). Labeling indicates treated cohorts that are significantly different from control in either longevity or steepness (quartile test), or in survival curves shape (AFT+KS test, results not shown). **b**, Volcano plots of the interventions presented in (**a**). On y-axis, p-values of the chi-square test described in Methods are plotted against, on x-axis, the combined effect sizes of longevity and steepness according to the following heuristic formula: $0.75*\ln(L_i/L_c)+0.25*\ln(S_i/S_c)$. Right panel is the zoom-in version of the left panel. Green, orange and purple symbols indicate male, female and mixed sex cohorts, respectively. The gray regions show that a threshold in effect size (0.2) is combined with a threshold in p-value (α=0.05 with Benjamini-Hochberg correction).

**Supplementary Data 2. Statistics of all analyzed interventions outside of the NIA ITP shown in Fig. 4b and Supp. Fig. 6.** Relative longevity, relative steepness, quartile test p-value and AFT+KS test statistic and p-value of all analyzed interventions outside of the NIA ITP.

**S7. Antioxidants NDGA and GTE increase steepness**

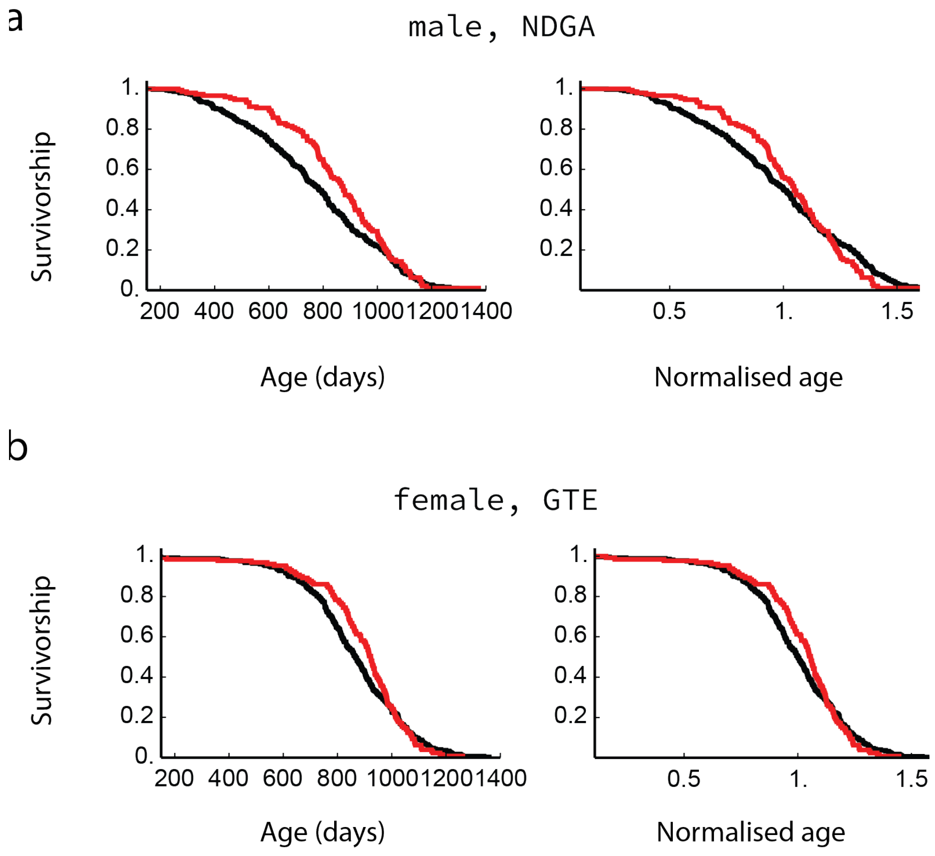


**Supplementary Figure 7. Survival curves and scaled survival curves for selected antioxidant supplements that steepens the survival curves. a**, NDGA (2500 ppm) in male mice starting at age of 9 months [57] (AFT+KS scaling test p=0.033) and **b**, GTE (2000 ppm) in female mice starting at age of 4 months [56] (AFT+KS scaling test p=0.023).

### S8. Rapamycin shows approximate scaling except at old ages

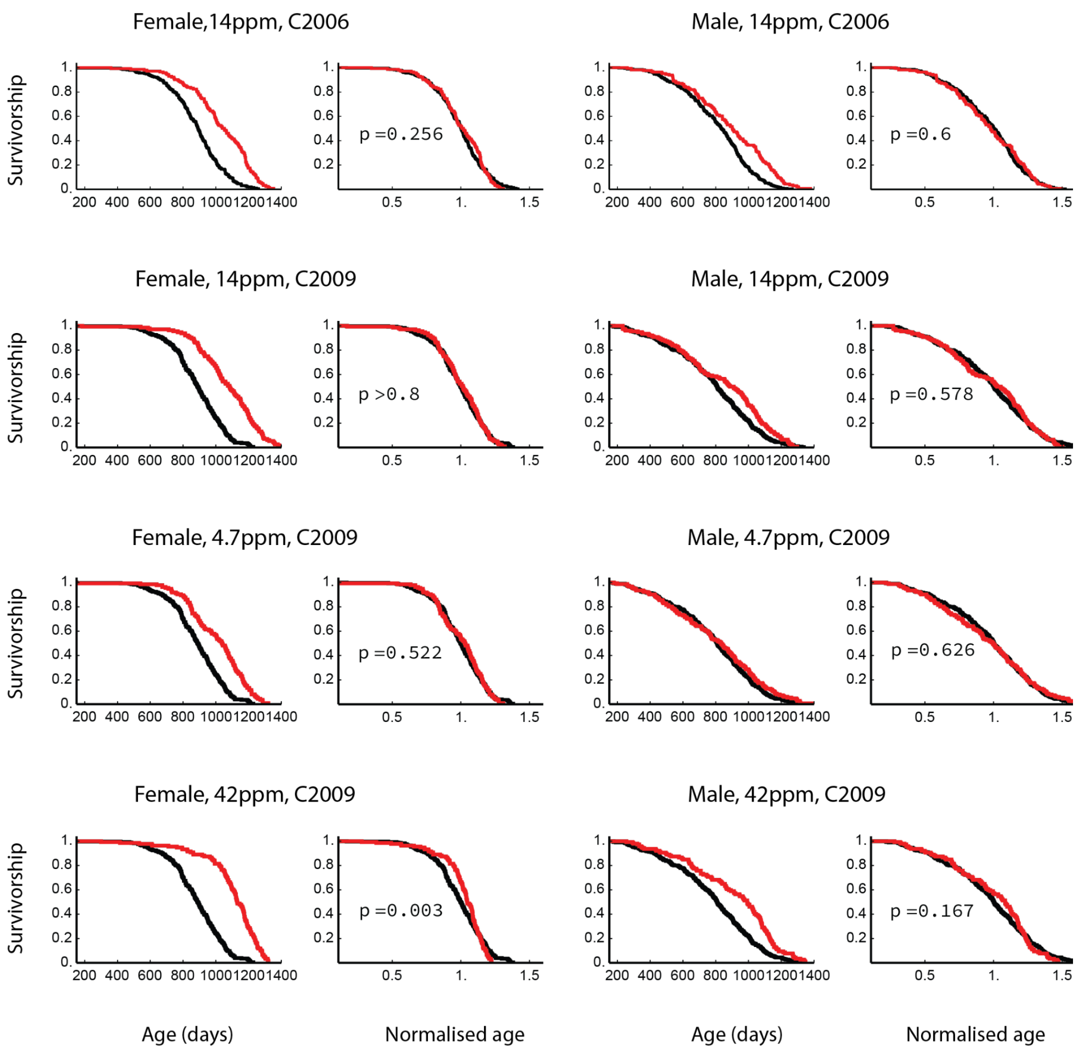


**Supplementary Figure 8. Survival curves and scaled survival curves for rapamycin [83,84] of female and male mice in the NIA dataset [50].** Rapamycin was at dose 42, 14 (twice) and 4.7 ppm starting from the age of 9 months. AFT+KS scaling test p-values are shown.

### S9. Rapamycin and acarbose interact additively to extend mouse lifespan

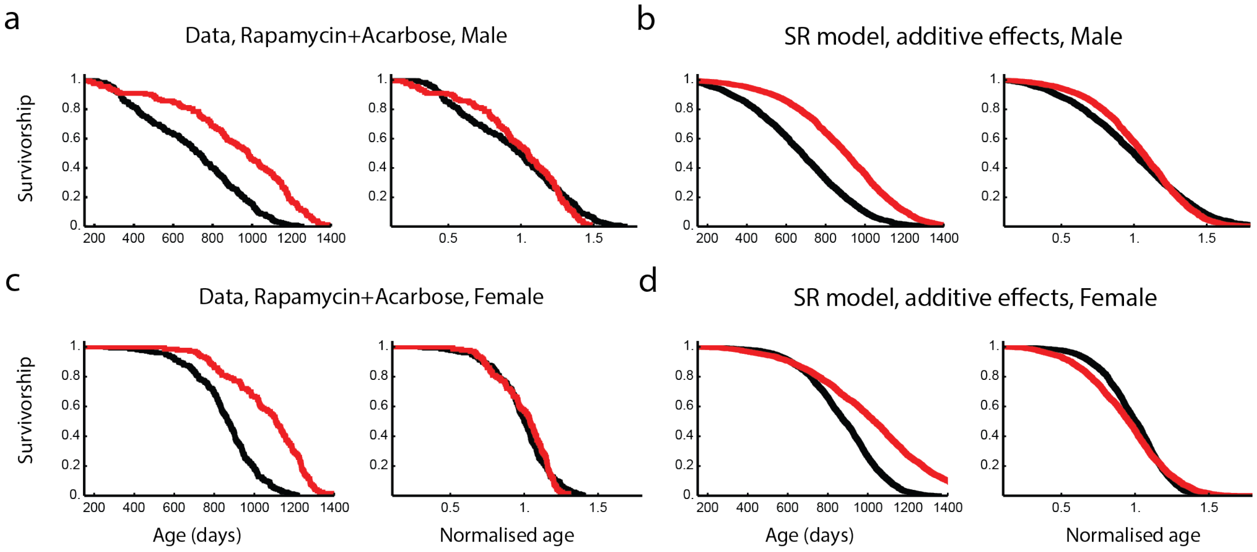


**Supplementary Figure 9. Rapamycin and acarbose affect the survival curves independently (non-epistatic) in mice. a**,**c,** Survival curves and scaled survival curves of, respectively, male and female mice treated by rapamycin+acarbose at 14.7ppm+1000ppm starting from the age of 9 months (red) and their controls (black). Data from NIA ITP C2017 cohorts. **b**,**d,** Simulated survival curves of the SR model and assuming the effects of the drugs are independent. which is to say that fold changes of parameters are simply added together to arrive at the model. Fold changes of SR model parameters $\eta$ and $\beta$ induced by the two drugs are estimated by matching the experimental longevity and steepness. Data used are rapamycin at 14ppm starting from the age of 9 months (C2006), and acarbose at 1000ppm starting from the age of 8 months (C2013). In all SR model simulations, $X_c$ is set to 23.2 and $\varepsilon$ is adjusted to 0.69 so that the simulations show longevity and steepness typical of experimental control cohorts. $\kappa$ is kept the same as in reference [32].

### S10. In vivo reprogramming shows approximate scaling in preliminary experimental data

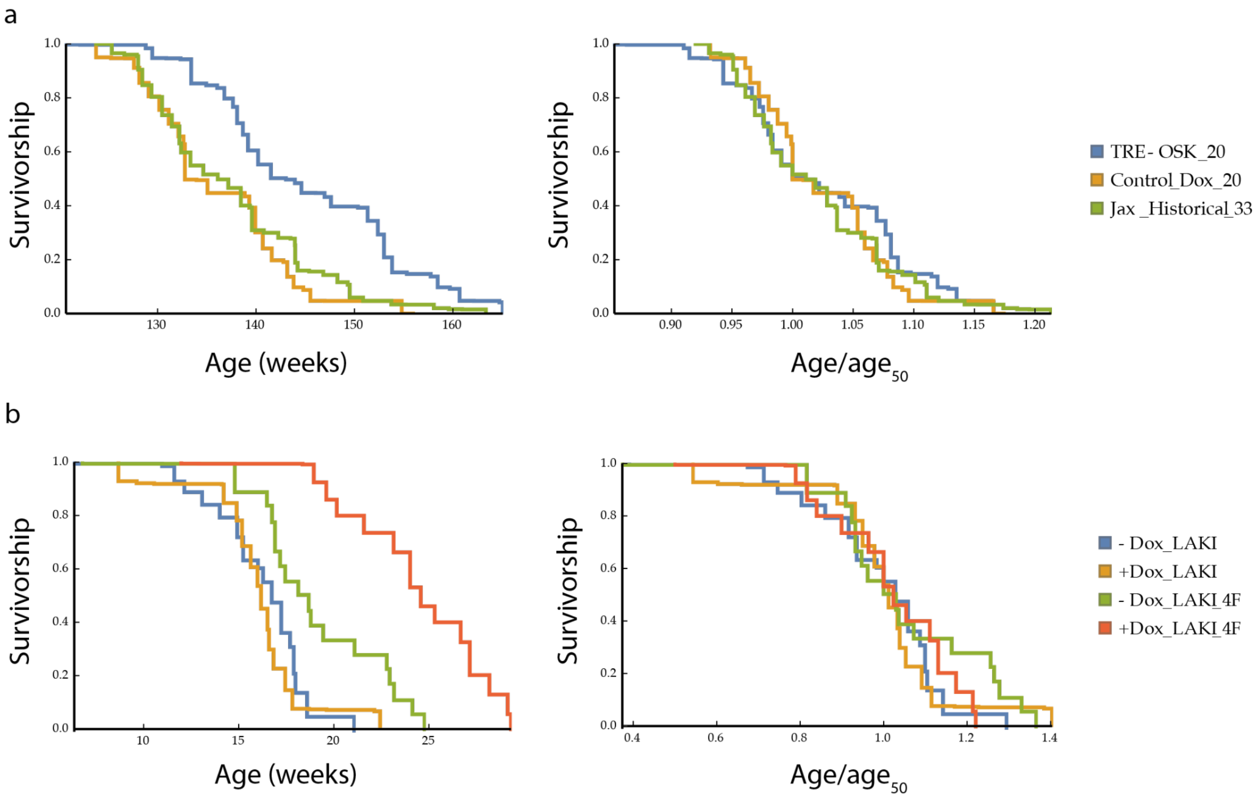


**Supplementary Figure 10.** ***In vivo*** **partial reprogramming survival curves and scaled survival curves.** **a**, Data from reference [24], injection is at week 124. **b**, Data from reference [23].
